# Mitigation of TDP-43-induced toxic phenotype by expression of RGNEF N-terminal fragment in ALS models

**DOI:** 10.1101/2023.09.29.560207

**Authors:** Cristian A. Droppelmann, Danae Campos-Melo, Veronica Noches, Crystal McLellan, Robert Szabla, Taylor A. Lyons, Hind Amzil, Benjamin Withers, Kirti S. Sonkar, Anne Simon, Emanuele Buratti, Murray Junop, Jamie M. Kramer, Michael J. Strong

**Author notes:** Corresponding Authors: Dr. Cristian Droppelmann and Dr. Michael J. Strong. These authors contributed equally to the work (Danae Campos-Melo, Veronica Noches).

## Abstract

Aggregation of the RNA-binding protein (RBP) TDP-43 is a hallmark of TDP-proteinopathies including amyotrophic lateral sclerosis (ALS) and frontotemporal dementia (FTD). Since TDP-43 aggregation and dysregulation are causative of neuronal death, there is a special interest in targeting this protein as a therapeutic approach. Previously, we found that TDP-43 extensively co-aggregated with the dual function protein (GEF (guanine exchange factor) and RBP) rho guanine nucleotide exchange factor (RGNEF) in ALS patients. Here, we show that a N-terminal fragment of RGNEF (NF242) interacts directly with the RNA recognition motifs (RRM) of TDP-43 competing with RNA, and that the IPT/TIG domain of NF242 is essential for this interaction. Genetical expression of NF242 in a fruit fly ALS model overexpressing TDP-43 suppressed the neuropathological phenotype increasing lifespan, abolishing motor defects, and preventing neurodegeneration. Intracerebroventricular injections of AAV9/NF242 in a severe TDP-43 murine model (rNLS8) improved lifespan and motor phenotype, and decreased neuroinflammation markers. Our results demonstrate an innovative way to target TDP-43 proteinopathies using a protein fragment with affinity for TDP-43, suggesting a promising therapeutic strategy for TDP-43 proteinopathies such as ALS and FTD.

## Introduction

Amyotrophic lateral sclerosis (ALS), also known as Lou Gehrig’s disease, is a neurodegenerative disorder characterized by progressive loss of voluntary muscle function, typically leading to death from respiratory failure within 3 to 5 years of symptom onset^1^. To date, despite multiple efforts, there is no effective therapy that arrests the progression of ALS because of the complex nature of its pathology^2^. The hallmark of ALS is the presence of unique protein inclusions, the most common of which are composed of RNA-binding proteins (RBPs)^3^ such as TAR DNA binding Protein (TDP-43), fused in sarcoma/translocated in liposarcoma (FUS/TLS), TATA-Box binding protein associated factor 15 (TAF 15), Ewing sarcoma breakpoint region 1 (EWS), RNA binding motif protein 45 (RBM45), heterogeneous nuclear ribonucleoprotein A1 and A2/B1 (hnRNPA1 and hnRNPA2B1), and rho guanine nucleotide exchange factor (RGNEF/p190RhoGEF)^4–13^. Of these proteins, TDP-43 is the most extensively studied ALS-associated RBP since its dysregulation has been directly associated with neuronal death *in vitro* and *in vivo*^14,15^ and TDP-43 immunoreactive neuronal cytoplasmic inclusions (NCIs) are observed in 97% of ALS cases^16^. TDP-43 is also the main pathological component of a group of diseases called TDP-43 proteinopathies which include frontotemporal dementia (FTD) and Limbic-Predominant Age-Related TDP-43 Encephalopathy (LATE)^17^. Because of this, there is a special interest in targeting TDP-43 as a therapeutic approach^18–20^.

Previously, we described that RGNEF forms extensive NCIs that co-aggregate with TDP-43 in motor neurons of ALS patients^11,12^ and observed that RGNEF works as a survival factor under stress conditions *in vitro*^21^. Also, we described that the amino-terminal fragment of RGNEF, called in this study NF242 (<u>N</u>H2-terminal <u>f</u>ragment of <u>242</u> amino acids), is part of a high molecular weight complex with TDP-43 *in vitro* and that both co-localize under metabolic stress conditions^22^.

Here, we hypothesized that NF242 works as a modifier of TDP-43 toxicity *in vivo*. To test this, we studied 1) the protein-protein interaction between NF242 and TDP-43 *in vitro* and *in silico*, 2) the co-expression of RGNEF or NF242 with TDP-43 in *Drosophila melanogaster*, and 3) the viral ectopic expression of NF242 in an aggressive murine model of ALS (rNLS8)^23^.

## Material and methods

### Antibodies, chemicals, and plasmids

Antibodies and other critical material used in this study are listed in the Supplementary Table 1.

### Constructs

To develop the transgenic flies, the coding regions of TDP-43^wt^, RGNEF, and flag-NF242 (previously described^22^) were cloned in the pTW-UASt vector (Drosophila Genomics Resource center), generating the pTW-TDP-43^wt^, pTW-RGNEF, and pTW-flag-NF242 vectors. For the luciferase reporter assay, the coding region of TDP-43^wt^ was cloned into the pcDNA-myc-His-A vector, generating the pcDNA-TDP-43-myc plasmid. For the complementation reporter assay (NanoBiT), the coding region of TDP-43^wt^, TDP-43-ΔNLS (nuclear localization signal of TDP-43 from amino acids 78 to 84 eliminated by site-directed mutagenesis), RGNEF, and NF242 were cloned into the pBiT1.1-C [TK LgBiT], pBiT1.1-N [TK LgBiT], pBiT2.1-C [TK SmBiT], and pBiT2.1-N [TK SmBiT] (Promega) vectors. pBiT constructs are detailed in the Supplementary Table 2. For SPR experiments the pQE30-TDP-43-RRM1 and pQE30-TDP-43-RRM2 plasmids used to express His-RRM-1 (amino acids 101 to 191 of TDP-43) and His-RRM-2 (amino acids 177 to 262 of TDP-43) were generated. The expression plasmid pQE30-TDP-43^1–296^ was used to express His-TDP-43^1–296^ (amino acids 1 to 261 of TDP-43, which include the amino terminal region and both RRMs). The expression plasmid pBAD-HisA-GST-TDP-43Cri was used to express His-GST-TDP-43^wt^. The expression plasmid pDEST566-RGNEF-275 was used to express His-MBP-RGNEF^1–275^.

### Cell lines

HEK293T cells (ATCC) were maintained in 25mM glucose, 1mM pyruvate Dulbecco’s modified Eagle’s medium (Gibco - Life technologies) containing 100U/ml penicillin, 100U/ml streptomycin (Gibco - Life technologies), 5µg/ml plasmocin (InvivoGen), and 10% fetal bovine serum (Gibco - Life technologies).

### Flies

Stocks and crosses of *Drosophila melanogaster* were cultured according to standard procedures and on standard fly food (water, yeast, cornmeal, brown sugar, agar, propionic acid, 10% methylparaben) (Bloomington Drosophila Stock Center). Flies were raised on 25°C and 70% humidity at a 12-hour day/night cycle.

*UAS-TDP-43^wt^*, *UAS-RGNEF*, and *UAS-flag-NF242* transgenic lines were generated by random germline insertion into *w1118* flies (*w^-^*) (BestGene). *GMR-Gal4*, *D42-Gal4*, and *elav-Gal4* driver lines were obtained from the Bloomington Drosophila Stock Center (Indiana University, Bloomington, Indiana, USA). The flies from stock centers used in this study are listed in the Supplementary Table 3.

Single transgenic flies homozygous for the transgene were used in the generation of the double transgenic fly lines, as well as in crosses with Gal4 drivers. Genotypes of the transgenic flies used in this study are listed in the Supplementary Table 4.

### Mice lines

Mice strains B6C3F1/J (JAX: 100010), B6;C3-Tg(NEFH-tTA)8Vle/J (JAX:025397), and B6;C3-Tg(tetO-TARDBP*)4Vle/J (JAX:014650) were purchased from The Jackson Laboratory. Experimental double transgenic mouse B6;C3-Tg(NEFH-tTA)8Vle Tg(tetO-TARDBP* (rNLS8) was generated after crossing JAX:025397 and JAX:014650. Double transgenic animals and the breeding pairs were maintained with doxycycline (50 μg/mL) in the drinking water to suppress the expression of TDP-43^23^. The wild type (wt) control mice for the experiments were obtained from the progeny of the crosses between JAX:025397 and JAX:014650 that were negative for both transgenes. For the experiments, rNLS8 males were excluded due to the observation of a urinary retention problem previously described for this transgenic line^24^.

### Study approval and animal housing

All procedures involving animals, surgeries, and animal maintenance were in accordance with the Canadian Council for Animal Care and the University Council on Animal Care guidelines for research. Ethics review and approval was granted by the Animal Care Committee of The University of Western Ontario (Protocol #2020-004). Mice were housed in the ACVS (Animal Care and Veterinary Services) in a temperature-controlled room (21– 23°C) with a 12 hr light–dark cycle. Animals were given free access to standard rodent chow and were provided with moistened chow on the cage floor and purified dietary supplement (Clear H2O DietGel 76A), after the Doxycycline was removed from the water for drinking.

### Transfections

Cell transfection of the constructs was performed using lipofectamine 2000 (Invitrogen) for the cytotoxicity assays or Magnetofection™ (OZ Biosciences) for the complementation reporter assay according to the manufacturer’s protocol.

### Complementation reporter assay (NanoBiT)

Protein-protein interaction was analyzed using the NanoBit Protein:Protein Interaction (PPI) System (Promega) according to the manufacturer’s instructions. Briefly, cells were seeded in white 96-well plates at 10,000 cells/ml per well and 24 hours after were transfected with the pBiT constructs listed in the Supplementary Table 1. After 48 hours, luciferase activity was measured using the Nano-Glo Live Cell Assay System (Promega) using a Luminometer (Modulus; Turner Biosystems).

### Protein Purification

His-GST-TDP-43^wt^, His-TDP-43^1–296^, His-RRM-1, His-RRM-2, and His-MBP-RGNEF^1–275^ recombinant proteins were purified from *E. coli* using the nickel-immobilized metal affinity chromatography (Ni-IMAC) method. For a detailed protocol see Supplementary Methods.

Purified His-TDP-43^1–102^ was generously provided by Dr. Stanley Dunn from the Department of Biochemistry at Western University (London, Canada). His-TDP-43^1–102^ contains an N-terminal 6xHistidine-Thioredoxin tag followed by the first 102 amino acids of human TDP-43.

### SDS-PAGE and western blot

To evaluate the purity of the purified proteins, SDS-PAGE and Western blot were performed (Supplementary Fig. 1). The protein aliquots from each purification were run in 4-20% Mini-PROTEAN® TGX™ Precast gradient gels. Gels were stained with Imperial™ Protein Stain (Thermo Scientific) or transferred to a nitrocellulose membrane. For the western blot the membrane was blocked in 5% BSA made in 1xTBST and the primary antibody (rabbit TDP-43) was incubated at 4°C with shaking overnight followed by HRP conjugated secondary antibodies for 60 minutes, at room temperature. Immunoblots were visualized using Western Lightning Plus Chemiluminescence Substrate (Perkin Elmer).

### Surface plasmon resonance (SPR) spectroscopy

Protein interactions were assessed using a Reichert 2SPR, SR7500DC System. Standard amine coupling (EDC/NHS chemistry) was used to capture purified His-MBP-RGNEF^1–275^ on a carboxymethyl dextran hydrogel sensor chip. The amount of ligand immobilized ranged from 2000-8000 µRIU. TDP-43 analyte proteins were serially diluted to the concentrations indicated in running buffer. His-GST-TDP-43^wt^ analysis was carried out using running buffer containing 20 mM Hepes-7.5, 50 mM KCl, 0.5 mM MgCl2, 50 mM NaCl, and 0.05% Tween-20. His-TDP-43^1–296^, His-RRM1, His-RRM-2, or His-TDP-43^1–102^ analysis were carried out in running buffer of 1X PBS and 0.1% Tween-20. In the experiments, 50-100 μl of TDP-43 analyte concentrations were injected on both the ligand and reference channels at 5-20 μl/min for 4-7 minutes with a 1-8-minute dissociation time at 22 °C. His-MBP protein (Supplementary Table 1) was used as control to evaluate possible unspecific binding. Dissociation constant (KD) analysis was performed using Reichert SPRAutolink (version 1.1.16), TraceDrawer (version 1.8.1), and GraphPad Prism 9.5 software packages.

### TDP-43-RGNEF interaction modeling

#### RGNEF domain analysis

The atomic coordinates of RGNEF residues 1-242 were extracted from the AlphaFold Protein structure database^25,26^ under the accession Q8N1W1. This model was queried for structural similarity against the entire PDB databank using the DALI protein structure comparison server^27^. The DALI results were analyzed using DALIview (https://github.com/rszabla/daliview) to reveal structurally similar domain families.

#### Structural prediction of the TDP-43 / RGNEF heterodimer

Constrained and unconstrained molecular docking of RGNEF^1–242^ onto TDP-43^96–269^ were performed using InterEvDock3 and ClusPro respectively^28,29^. The atomic coordinates of RGNEF^1–242^ were taken from the AlphaFold structure database while those of TDP-43 were taken from the available NMR structure^30^. To generate the dimer model, the sequence of RGNEF^1–242^ (Uniprot accession Q8N1W1) and the sequence of full-length TDP-43 (Uniprot Q13148) were both used as inputs for AlphaFold2 running in complex prediction mode on CollabFold^26,31^.

The top-scoring output model was used for further structural minimizations. For this, the structure of TDP-43 in the dimer structure was limited to the two RRM domains with about 8 additional flanking residues on either side (residues 96-269)^32^. The top-scoring model was used for residue-contact analysis.

The relative structural stability of the dimer was quantified by measuring the conformational spread between all output models. This was done using a custom PyMOL script that aligned each of the 1,000 output models against the top-scoring model for the dimer and calculating an RMSD value for each model. The top 50 scoring models for the dimer were deposited to the ModelArchive database as a multi-model PDB file.

TDP-43(96–269) bound to a 12-mer strand of RNA was also minimized from experimental NMR coordinates^30^ (PDB accession: 4BS2).

### *NEFL* mRNA stabilization activity

*NEFL* 3’UTR stability by TDP-43 (pCDNA-TDP-43-myc) was studied using a luciferase reporter assay as previously described^12^ with minor modifications (pcDNA-flag-NF242 plasmid^22^ was used for the co-expression of flag-NF242).

### Cytotoxicity analysis

Cells were seeded in white 96-well plates at 9,000 cells/ml per well. Cytotoxicity was measured using the CytoTox-Glo™ Cytotoxicity Assay kit (Promega) according to the manufacturer’s protocol after 2 days of transfection. To obtain the percentage of cell toxicity, the values obtained after the stress condition or control were normalized against total protease activity obtained after cell lysis using digitonin.

### Expression analysis in flies

To check the expression of the TDP-43, RGNEF, NF242, or GFP in our fly models, total RNA from at least 15 flies was isolated using Trizol reagent (Invitrogen). Reverse transcription was performed using the Superscript II reverse transcriptase system (Invitrogen). PCR reactions (Supplementary Fig. 2) were performed using primers listed in the

Supplementary Table 5.

### Lifespan analysis in flies

F1 male progeny of transgenic flies *elav>RGNEF*, *elav>RGNEF;TDP-43*, *elav>NF242;TDP-43*, *elav>GFP;TDP-43*, *D42>RGNEF;TDP-43*, *D42>NF242;TDP-43*, and *D42>GFP;TDP-43* were collected and maintained in vials in an incubator set to 25°C at 70% humidity with controlled day/night cycles. The number of dead and live flies were counted every other day. Heterozygote driver lines elav>*w^-^*, *D42>w^-^*, and non-expressing *UAS-RGNEF* flies were used as additional controls.

### Motor analysis in flies

F1 male progeny of transgenic flies *elav>RGNEF*, *elav>RGNEF;TDP-43*, *elav>NF242;TDP-43*, *elav>GFP;TDP-43*, *D42>RGNEF;TDP-43*, *D42>NF242;TDP-43*, and *D42>GFP;TDP-43* were collected to evaluate the negative geotaxis (locomotion) using a climbing assay. To do this, flies were transferred to a graduated cylinder (24 cm height, 3 cm of diameter) divided into 4 vertical quadrants (from the lower part: quadrant 1 to 3 of 5 cm each, and quadrant 4 of 9 cm) and sealed with parafilm. Flies were tapped to the bottom of the cylinder and the number of flies present in each quadrant was recorded at 10s and 20s. Measurements were repeated a total of 4 times every 3 days. Climbing index was calculated using the formula: Climbing index = [Q1 + (Q2 *x* 2) + (Q3 *x* 3) + (Q4 *x* 4)] / Total number of flies where Q represents the number of flies in the respective quadrant^33^.

### Fly eye degeneration

F1 male progeny of transgenic flies *GMR>RGNEF*, *GMR>NF242*, *GMR>RGNEF;TDP-43*, *GMR>NF242;TDP-43, GMR>GFP;TDP-43*, and *GMR>w^-^* were collected to capture images of fly eyes. Flies were anaesthetized with CO2 and then photographed using a Leica S9i Stereomicroscope (Leica Microsystems Inc.).

### Fixation of fly tissues

The fixation protocol from the Shcherbata group^34^ was performed to obtain paraffin-embedded adult flies. Whole flies in collars were first incubated in Carnoy’s solution containing absolute ethanol, chloroform, and glacial acetic acid 6:3:1 ratio, overnight at 4°C. Flies were then dehydrated by incubation in 40% ethanol for 20 minutes, 70% ethanol for 20 minutes and twice in 100% ethanol for 10 minutes each. Afterwards, flies were incubated in methylbenzoate (MB) and MB with paraffin solution, 1:1 ratio, for 30 minutes each at 60°C following which they were incubated twice in paraffin solution for 60 minutes each at 60°C. Flies in paraffin were then allowed to solidify at room temperature overnight before cutting the paraffin-embedded flies into 7 μm sections (Pathology Core Facility, Robarts Research Institute). Hematoxylin-eosin staining of selected slides was performed for checking quality and anatomy visualization.

### Immunofluorescence for flies and mice

For slide deparaffinizing, sections of fly brain and eye tissue, mice brain, or mice spinal cord were first seated on a slide warmer at 60°C for 30 minutes. Then slides were rehydrated in a series of graded alcohols and water. Antigen retrieval was performed in a pressure cooker for 30 minutes at 100°C in a buffer containing 10mM citric acid, 2mM EDTA and 0.05% Tween-20 pH 6.2 for fly tissues or in 10mM sodium citrate, 0.05% Tween, pH 6.0 for mouse tissues. Next, slides were incubated for 60 minutes at room temperature in PBS pH 7.2 blocking solution with 5% BSA and 0.3% Triton-X 100, and with primary antibodies at 4°C overnight in a humidifying chamber. After the washes, slides were incubated with Alexa Fluor secondary antibodies for 60 minutes at room temperature. Dilutions for primary and secondary antibodies are indicated in Supplementary Table 6. Slides were then incubated with 2μg/ml Hoechst for 3 minutes. After the washes and once dry, coverslips were mounted to the slides using a fluorescent mounting media (Dako). Slides were examined using an SP8 Lightening Confocal microscopy system (Leica Microsystems Inc.). For the super-resolution stimulated emission depletion (STED) microscopy images a Leica STELLARIS STED microscope (Leica Microsystems Inc.) was used. The multi-STED method was performed using Alexa-488 and Alexa-595 fluorophores and the 592 nm and 775 nm STED depletion lasers for tau-STED analysis. All images were visualized using the LAS X 2.0 software (Leica Microsystems Inc.).

### Co-localization images

Intensity Correlation Analysis^35^ using ImageJ software was performed to obtain the co-localization images. Co-localized pixels are shown as PDM (Product of the Differences from the Mean) images. PDM=(red intensity-mean red intensity)×(green intensity-mean green intensity). In the co-localization images, blue and purple colors indicate lower level of co-localization while yellow and white indicate a high level of co-localization.

### Intracerebroventricular injections of AAVs

Self-complementary adeno-associated viruses serotype 9 for neuronal-specific expression of GFP (AAV9/GFP) and NF242 (AAV9/NF242) were produced to a yield of 2.0×10^13^ GC/ml and 2.4×10^13^ GC/ml respectively (Vector Biolabs) using pscAAV-GFP and pscAAV-NF242 plasmids^22^. Mice were stereotaxically injected with the AAVs intraventricularly in the brain (injection site: AP = -0.4 mm; ML = -1.0, +1.0 mm; and DV = 2.3 mm from Bregma) with 2.5 μL (bilateral) of AAV9/GFP or AAV9/NF242 at a rate of 1 μl/min with a 33-G Hamilton syringe. One week after the surgery doxycycline was removed from the double transgenic mice to induce the expression of TDP-43-ΔNLS.

### Motor analysis in mice

Motor tests for the mice were performed once per week from doxycycline retrieval.

#### Clasping

Mice were suspended by the tail ∼30 cm above the cage and slowly lowered. Clasping of both hindlimbs that was maintained for ∼30 s was recorded as a positive response^36^.

#### Grip-strength assessment

Front limbs strength was assessed using a Model Grip Strength meter (Columbus Instruments) horizontally mounted. Mice had to grip a wire bail attached to a force transducer sensing shaft (Chatillon 2LBF AMETEK). The peak force of 5 trials was the grip strength expressed in normalized force (Newton/grams)^37,38^.

#### Rotarod

To test motor coordination and balance^37^ mice were placed on a rotarod apparatus (AccuRotor Rota-Rod, Omnithech electronics, Inc.; software, Fusion 6.4 AccuRotor edition) at a speed of 4 rpm with increased linearly acceleration up to 40 rpm over 300s. After the initial training session, weekly session of 4 trials were performed for each animal and the average in the latency to fall of 4 trials was calculated.

#### Catwalk

CatWalk XT® Version 10.6 system by Noldus (Leesburg, VA, USA) was used for mice gait assessment. Tests were conducted in a room with red light and the analysis was made with the average of two videos per animal. Runs were analyzed using Noldus software^39,40^.

#### OpenField

Mice were placed in a square arena (20cm x 20cm) (AccuScan Instruments Inc. Columbus, Ohio USA) and activity for 20 min was recorded by infra-red photo beams along x, y, z axes using software Fusion V5 VersaMax Edition. Distance travelled (converted from beam breaks to cm) was recorded at 5-min blocks^41^. The results of open field were not compared with their wt counterpart because of the hypermobility associated with TDP-43 transgenic models^42,43^, which creates a different basal for transgenic mice when compared with wt mice.

### Mice end point

Disease end stage in mice was defined as: CS 4 (clinical score; functional paralysis of both hindlimbs), CS 4+ (CS 4 plus loss of body weight > 20% or body condition score <2), and CS 5 (CS 4 plus righting reflex >20 s)^44^.

### Pathology quantification

Relative fluorescence intensity of TDP-43, GFAP and Iba1 staining in the ventral horn of the lumbar spinal cord or brain cortex of rNLS8 mice injected with AAV9/GFP and AAV9/NF242 was measured using LAS X 2.0 software (Leica), quantifying the intensity on at least 5 different slices (technical replicates) for each animal.

### Statistical analysis

The statistical analyses were performed with GraphPad Prism 9.5 software. Log-rank (Mantel-Cox) test analysis was used to compare lifespan curves and clasping. For the protein-protein experiments, one-way ANOVA with Dunnett’s post-hoc or Student’s t-test were performed. For the animal motor test studies two-way ANOVA analysis comparing the difference between treatments was performed. For the pathology quantification Student’s t-test were performed. Data were expressed as mean ± SEM. Data was judged to be statistically significant when p < 0.05

## Results

### Interaction between NF242 and TDP-43

Previously, we observed that RGNEF and TDP-43 colocalize and co-immunoprecipitate and that NF242, an amino-terminal fragment of RGNEF encompassing its first 242 amino acids, and TDP-43 are part of a high molecular complex^12,22^. To evaluate if the interaction between RGNEF with TDP-43 is direct, we performed a complementation reporter assay (NanoBiT)^45^ in HEK293T cells. We transfected a series of constructs containing RGNEF, NF242, TDP-43^ΔNLS^, and TDP-43^wt^ fused to the large or small subunit of the luciferase (Supplementary Fig. 3a-b). TDP-43^ΔNLS^ has been previously described to emulate pathological conditions and localizes in the cytoplasm^46^ which we thought would facilitate the interaction with RGNEF (mainly cytoplasmic^12,22^). The amino- or carboxy-terminal end of TDP-43^ΔNLS^ fused to the large subunit of luciferase showed interaction with both RGNEF and NF242, but only when the carboxy-terminal end of the latter proteins was fused to the small subunit of the luciferase (Supplementary Fig. 3c and Supplementary Fig. 4a-d). In experiments with TDP-43^wt^, we observed interaction only with NF242 and when both proteins had the luciferase subunit fused to the carboxy-terminal end (Supplementary Fig. 3c and Supplementary Fig. 4e-h).

To further validate the interaction between NF242 and TDP-43, we measured complex formation between the two proteins directly via surface plasmon resonance spectroscopy (SPR). In this assay, NF242 was fixed to the SPR substrate as the immobilized ligand and TDP-43 was injected as the mobile analyte. To obtain sufficient amounts of purified protein for the SPR assay, we used optimized constructs of NF242 and TDP-43 which maximize recombinant expression and purification efficiency in *E. coli*. For TDP-43, we expressed the full-length wild-type protein with an N-terminal GST fusion (His-GST-TDP-43^wt^). For NF242, we included an N-terminal MBP fusion and extended the C-terminal truncation boundary of N242 by 33 residues (His-MBP-RGNEF^1–275^). The SPR experiments showed direct interaction between the proteins (Fig. 1a) with a KD of 1.78 ± 0.49 μM (n=4). Then, we evaluated which region of TDP-43 was critical for the interaction with NF242. For NanoBiT assays, we used a series of TDP-43 constructs with deletions of different domains of the protein with the luciferase subunits fused to the carboxy-terminal end of both TDP-43 and NF242. We observed that the constructs lacking the carboxy-terminal region of TDP-43 (TDP-43^1–366^ and TDP-43^1–274^) maintained the interaction with NF242 (Fig. 1b-d). However, when the RNA recognition motifs (RRMs) domains were removed (TDP-43^ΔRRM1–2^), no interaction was observed (Fig. 1e). SPR experiments using His-TDP-43^1–296^ as analyte confirmed the importance of the amino-terminal region of TDP-43 (including both RRMs) for the direct interaction with NF242 (Fig. 1f) with a KD of 4.11 ± 1.33 μM (n=3). When the protein His-TDP-43^1–102^ which lacks both RRMs was used as analyte, no interaction was detected between the proteins (Supplementary Fig. 4i).

**Fig. 1.**
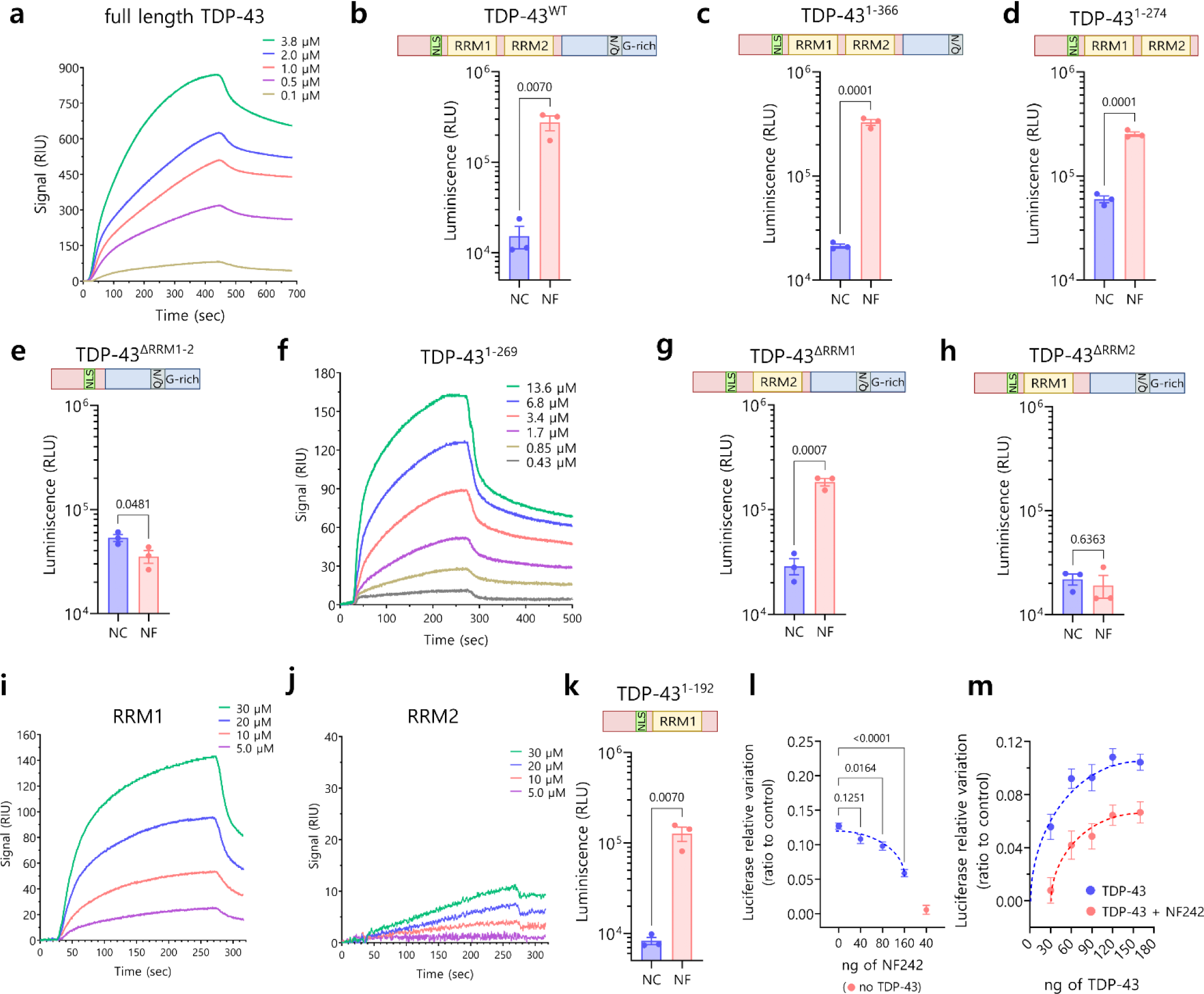
Interaction between RGNEF/NF242 and TDP-43. a, Representative SPR sensorgrams showing the interaction between His-GST-TDP-43^wt^ (analyte) and His-MBP-RGNEF^1–275^ (ligand) at different concentrations of His-GST-TDP-43^wt^. KD = 1.78 ± 0.49 μM (n=4). b, NanoBiT experiment showing interaction between TDP-43^wt^ (structure detailed) and NF242 (NF) (n=3; NC= negative control). c, NanoBiT experiment showing interaction between TDP-43^1–366^ (structure detailed) and NF242 (NF) (n=3; NC= negative control). d, NanoBiT experiment showing interaction between TDP-43^1–274^ (structure detailed) and NF242 (NF) (n=3; NC= negative control). e, NanoBiT experiment showing an absence of interaction between TDP-43^ΔRRM1–2^ (structure detailed) and NF242 (NF) (n=3; NC= negative control). f, Representative SPR sensorgrams showing the interaction between His-TDP-43^1–296^ (analyte) and His-MBP-RGNEF^1–275^ (ligand) at different concentrations of His-TDP-43^1–296^. KD = 4.11 ± 1.33 μM (n=3). g, NanoBiT experiment showing interaction between TDP-43^ΔRRM1^ (structure detailed) and NF242 (NF) (n=3; NC= negative control). h, NanoBiT experiment showing the lack of interaction between TDP-43^ΔRRM2^ (structure detailed) and NF242 (NF) (n=3; NC= negative control). i, Representative SPR sensorgrams demonstrating the interaction between His-RRM1 (analyte) and His-MBP-RGNEF^1–275^ (ligand) at different concentrations of His-RRM1(n=4). j, Representative SPR sensorgrams demonstrating weak interaction (low signal intensity) between His-RRM2 (analyte) and His-MBP-RGNEF^1–275^ (ligand) at different concentrations of His-RRM2 (n=4). k, NanoBiT experiment showing interaction between TDP-43^1–192^ (structure detailed) and NF242 (NF) (n=3; NC= negative control). l, Quantification of TDP-43 stabilizing activity over *NEFL* 3’UTR (fixed amount of TDP-43) in presence of increasing amounts of NF242 (blue dots, n=3). NF242 decreases TDP-43 stabilizing activity in a dose-dependent manner. Red dot shows the control in absence of TDP-43 (n=3). m, Quantification of TDP-43 stabilizing activity over *NEFL* 3’UTR in presence (red) of absence (blue) of 120 ng of NF242 at increasing amounts of TDP-43. Displacement of the dose-response curve suggests competition of NF242 and RNA for TDP-43 (n=3).

To evaluate which RRM of TDP-43 was critical for the interaction with NF242, we created two NanoBiT constructs of TDP-43 lacking RRM1 (TDP-43^ΔRRM1^) or RRM2 (TDP-43^ΔRRM2^). The deletion of RRM1 did not alter the interaction between TDP-43 and NF242 (Fig. 1g), but the deletion of RRM2 completely abolished it (Fig. 1h). SPR using His-RRM1 and His-RRM2 as ligands showed robust interaction between His-RRM1 and His-MBP-RGNEF^1–275^ (Fig. 1i) but weak interaction with His-RRM2 (Fig. 1j). To reconcile the NanoBiT and SPR results about the role of RRM1/2 in TDP-43-NF242 interaction, we evaluated if the carboxy-terminal region of TDP-43^ΔRRM2^ was blocking the access of NF242 to TDP-43. We generated a TDP-43^1–192^ NanoBiT construct which lacks RRM2 and the carboxy-terminal domain of the protein. Our analysis showed an interaction between TDP-43^1–192^ and NF242, confirming the blocking effect by the carboxy-terminal domain of TDP-43 (Fig. 1k). These results suggest that both RRMs are necessary for the interaction between TDP-43 and NF242 and that RRM1 is the domain that has the strongest interaction with NF242.

To study whether the interaction with NF242 had any functional consequence for TDP-43, we used a luciferase assay that we previously developed to evaluate the regulation of the stability of *NEFL* 3’UTR by TDP-43^12^. We observed a moderate inhibition of NF242 over the RNA stabilizing activity of TDP-43 suggesting a competitive effect between NF242 and RNA for TDP-43 (Fig. 1l-m).

Molecular docking modeling also demonstrated the importance of both RRMs of TDP-43 in the interaction with NF242 and predicted that NF242 binds to the RNA binding site of TDP-43 (Fig. 2a-d). *In silico* analysis suggested that the amino acids 76 to 81, corresponding to a loop in the IPT/TIG (immunoglobulin, plexins, transcription factors-like/transcription factor immunoglobulin) domain (amino acids 1 to 95) of NF242 (Supplementary Fig. 3a), are critical for the interaction with TDP-43 in the interface between the RRM1 and RRM2 domains (Fig. 2e-f). To test this, we generated two NanoBiT constructs with mutations in the loop region (Fig. 2g). We observed no interaction between NF242 and TDP-43 when the loop 76-81 of NF242 was disrupted (Fig. 2h).

**Fig. 2.**
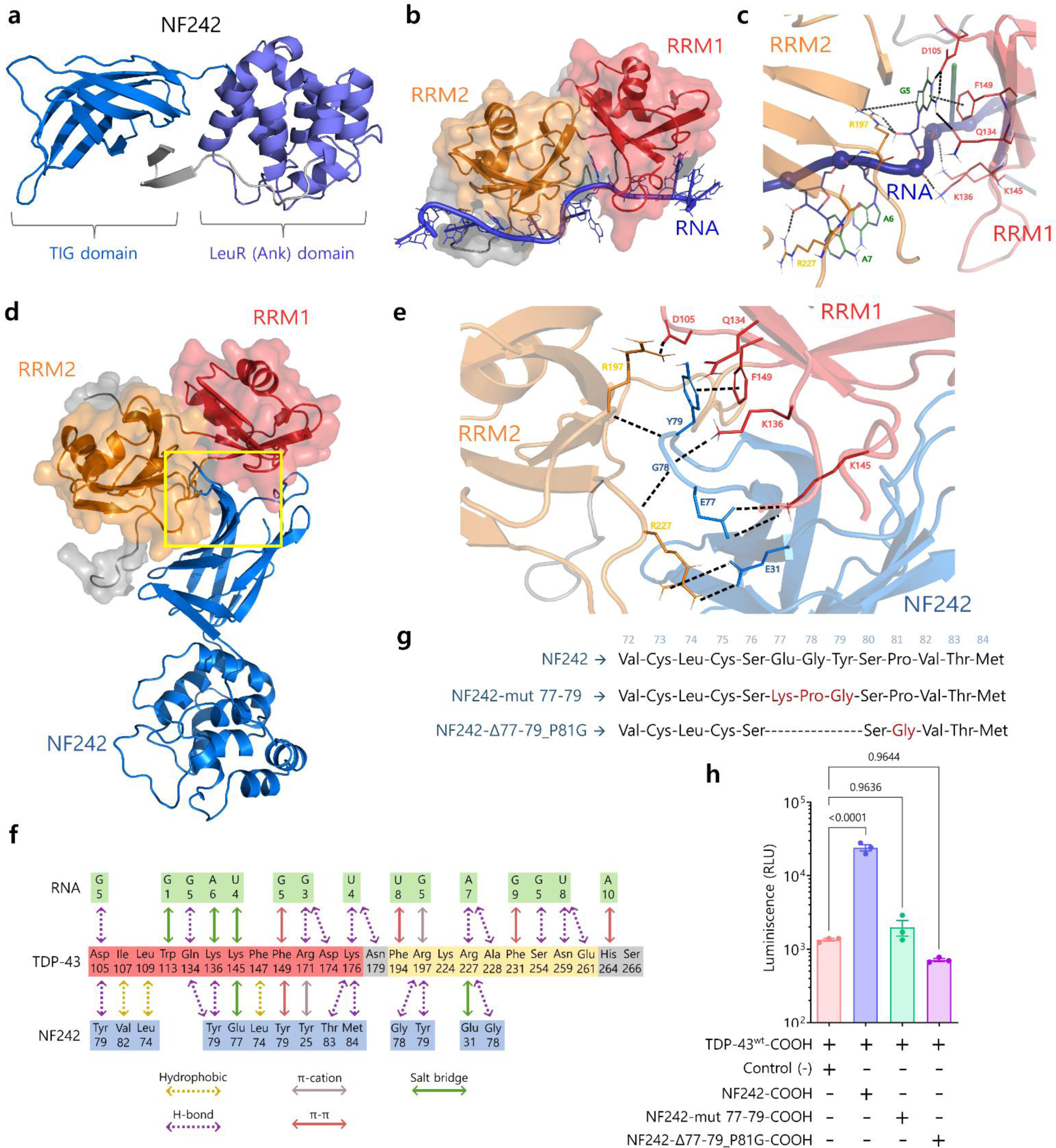
Modeling of TDP-43-NF242 interaction. a, NF242 structure based in the atomic coordinates of RGNEF residues 1-242 (NF242) extracted from the AlphaFold Protein structure database (accession Q8N1W1). b, Minimized structure of TDP-43 in complex with AUG12 RNA from experimental NMR coordinates (PDB accession: 4BS2). c, Region of high inter-molecular contacts occurring between TDP-43 and RNA. d, Minimized structure of TDP-43 in the complex with NF242. e, Region of high inter-molecular contacts occurring between TDP-43 and NF242 (yellow square in d) showing the most important amino acids interactions from the loop 76-81 of NF242 and the interface between RRM1 and RRM2 of TDP-43. f, Summary of all intermolecular contacts. g, Schematic showing the mutants used to study the importance of the loop 76-81 of the TIG domain of NF242 in the interaction with TDP-43. h, NanoBiT experiment showing that the mutants NF242-mut 77-79 and NF242-Δ77-79_P81G, both fused to smBiT in the carboxy-terminal end, do not interact with TDP-43 (p=0.9636 and p=0.9644 respectively). The interaction with NF242 is shown as positive control (p<0.0001).

These results suggest that the interaction between NF242 and TDP-43 could have consequences *in vivo* in ALS models.

### NF242 and TDP-43 co-expression in flies

After previously determining that RGNEF has a protective effect in cells under stress^21^, we sought to evaluate whether RGNEF exerts this protection when TDP-43 is overexpressed. We co-transfected HEK293T cells with plasmids expressing RGNEF and TDP-43^wt^ and observed that RGNEF decreased TDP-43-induced cytotoxicity compared to TDP-43^wt^ overexpression alone (Supplementary Fig. 5a). The same result was obtained when NF242 and TDP-43^wt^ were co-expressed (Supplementary Fig. 5b). This observation and the direct interaction between NF242 and TDP-43 led us to study the *in vivo* effect of the co-expression of either NF242 or RGNEF with TDP-43^wt^. We created lines of transgenic *Drosophila melanogaster* (fruit flies) co-expressing RGNEF and TDP-43^wt^ or NF242 and TDP-43^wt^ using the UAS-GAL4 system^47^. As neuropathological TDP-43 positive control for the experiment, we created the *GFP;TDP-43^wt^* fly, which incorporated GFP under the UAS promoter to compare only double transgenic flies and account for any possible effect caused by GAL4 acting over two UAS promoters (Supplementary Fig. 6).

First, we analyzed the effect of the expression of RGNEF alone on the lifespan of the flies. When RGNEF was overexpressed using the elav pan-neuronal driver (*elav>RGNEF* line), we observed an increased lifespan of the flies (average of 72.89 ± 1.22 days) compared with the heterozygous driver alone control *elav>w^-^* (*elav* crossed with the *w-* line) (average of 54.95 ± 0.88 days; p<0.0001; w^-^ is the parental line for the transgenic flies) and the *RGNEF* line without the driver (average of 56.24 ± 1.01 days; p=0.0035; Fig. 3a). Then, we evaluated the effect of the co-expression of RGNEF or NF242 with TDP-43 (*elav>RGNEF;TDP-43^wt^* and *elav>NF242;TDP-43^wt^* lines) on the flies’ lifespan. When compared with the *elav>GFP;TDP-43^wt^* line which had a short lifespan (average of 4.27 ± 0.13 days) consistent with previous reports^48–50^, both *elav>RGNEF;TDP-43^wt^* and *elav>NF242;TDP-43^wt^* lines showed a significantly longer lifespan (average of 63.73 ± 2.25 and 69.56 ± 1.44 days respectively; p<0.0001; Fig. 3b). We obtained similar results using the D42 motor neuron driver; *D42>GFP;TDP-43^wt^* line had a significantly shorter lifespan (average 14.13 ± 0.31 days) than *D42>NF242;TDP-43^wt^* line and the heterozygous driver alone control *D42>w^-^* line (average 61.07 ± 1.06 and 53.32 ± 1.44 days respectively; p<0.0001; Fig. 3c).

**Fig. 3.**
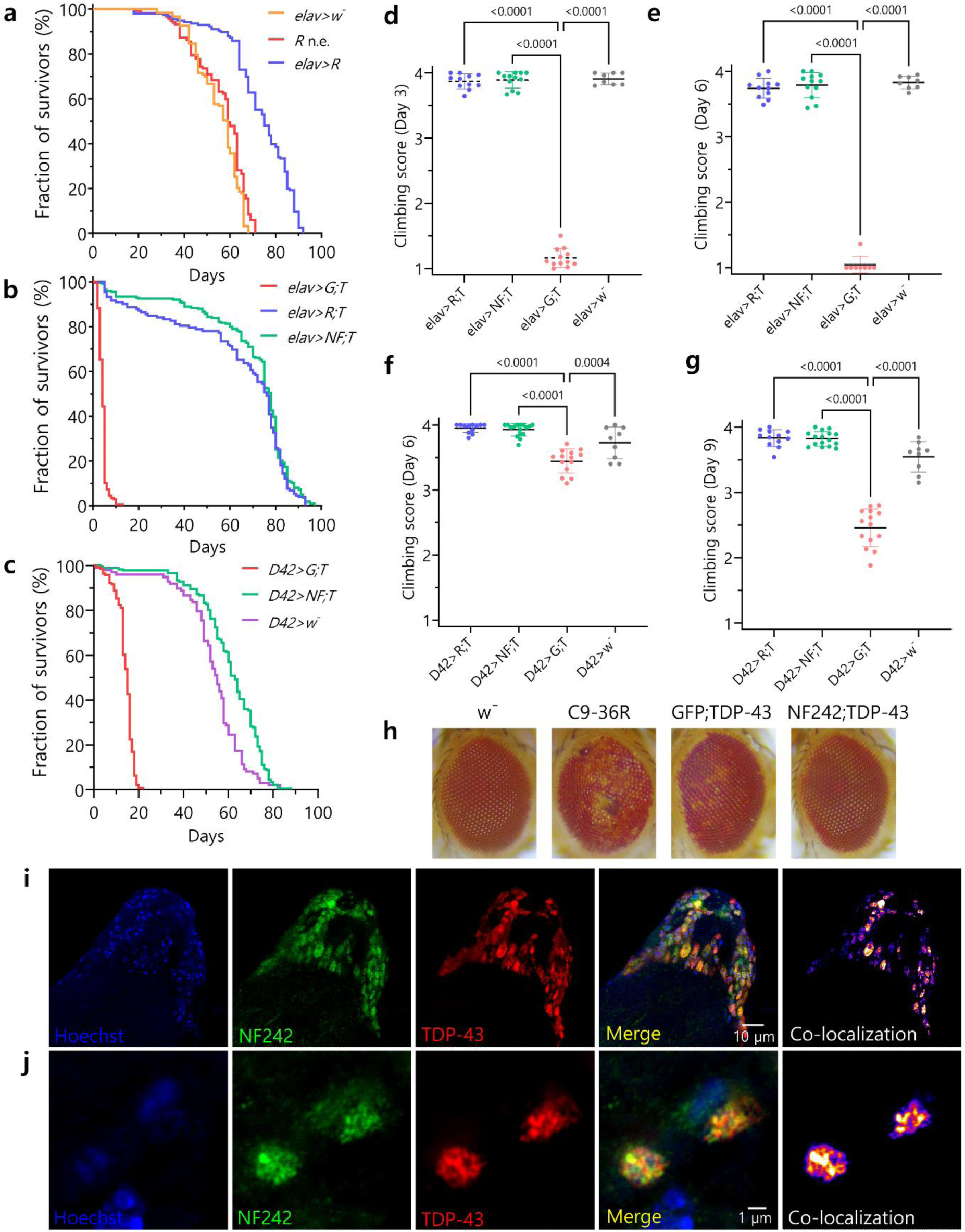
Co-expression of RGNEF or NF242 with TDP-43 in fruit flies. a, Kaplan-Meier graph showing the survival of elav>RGNEF (elav>R; n=156), RGNEF no driver control (R; n=117), and elav>w-(control of driver crossed with parental line; n=123). The elav>RGNEF line shows an increased lifespan compared to RGNEF no driver control (p=0.0035) and elav>w-(p<0.0001) lines. b, Kaplan-Meier graph showing the survival of *elav>GFP;TDP-43^wt^* (*elav>G;T*; n=178), *elav>RGNEF;TDP-43^wt^* (*elav>R;T*, n=132), and *elav>NF242;TDP-43^wt^* (*elav>NF;T*, n=224). The *elav>GFP;TDP-43^wt^* line shows a reduced lifespan, an effect that is suppressed in the *elav>RGNEF; TDP-43^wt^* (p<0.0001) and *elav>NF242;TDP-43^wt^* (p<0.0001) lines. c, Kaplan-Meier graph showing the survival of *D42>GFP; TDP-43^wt^* (*D42>G;T*; n=143), *D42>NF242;TDP-43^wt^* (*D42>NF;T*, n=181), and *D42>w^-^* (control of driver crossed with parental line, n=98) . The *D42>GFP;TDP-43^wt^* line shows a reduced lifespan, an effect that is suppressed in the *D42>NF242* line (p<0.0001). The latter also show an increase in lifespan compared to the control *D42>w^-^* line (p<0.0001). d-e, Negative geotaxis assay showing the climbing score at day 3 and 6 for *elav>RGNEF;TDP-43^wt^* (*elav>R;T*, n=11; 110 flies), *elav>NF242;TDP-43^wt^* (*elav>NF;T*, n=12, 120 flies), *elav>GFP;TDP-43^wt^* (*elav>G;T*; with n=12; 120 flies at day 1), and *elav>w^-^* (n=8; 80 flies) lines. The *elav>GFP;TDP-43^wt^* line shows a severe motor phenotype that is suppressed when RGNEF or NF242 is co-expressed with TDP-43^wt^ in neurons (p<0.0001). f-g, Negative geotaxis assay showing the climbing score at day 6 and 9 for *D42>RGNEF;TDP-43^wt^* (*D42>R;T*, n=12; 120 flies), and *D42>NF242;TDP-43^wt^ (D42>NF;T*, n=16, 160 flies), *D42>GFP;TDP-43^wt^* (*D42>G;T*; n=14; 140 flies), and *D42>w^-^* (n=9; 90 flies). The *D42>GFP; TDP-43^wt^* line shows a significant motor phenotype that is suppressed when RGNEF or NF242 is co-expressed with TDP-43^wt^ in motor neurons (p<0.0001). h, Representative images showing the eye phenotype of *GMR>w^-^* (negative control), *GMR>36R* (positive control), *GMR>GFP;TDP-43^wt^*, and *GMR>NF242;TDP-43^wt^* lines. NF242 co-expression with TDP-43^wt^ suppresses the eye degeneration observed in the *GMR>NF242;TDP-43^wt^* line. i, Immunofluorescence of adult *elav>NF242;TDP-43^wt^* fly brain tissue showing the co-localization between NF242 and TDP-43^wt^ in neurons. j, Confocal image at higher magnification of adult *elav>NF242;TDP-43^wt^* fly brain tissue showing the co-aggregation between NF242 and TDP-43^wt^ in neurons.

The effect of RGNEF and NF242 expression in the motor phenotype induced by TDP-43^wt^ in flies was evaluated using a negative geotaxis assay^51,52^. We observed that the toxic motor phenotype induced by TDP-43^wt^ under the neuron-specific elav driver (*elav>GFP;TDP-43^wt^* line) was suppressed by either RGNEF (*elav>RGNEF;TDP-43^wt^* line) or NF242 (*elav>NF242;TDP-43^wt^* line; Fig. 3d-e). Analogous results were observed when these proteins were expressed only in motor neurons using the D42 driver (Fig. 3f-g). The effect of TDP-43^wt^, RGNEF and NF242 expression in the induction of eye degeneration was studied using the eye specific GMR driver. *GMR>GFP;TDP-43^wt^* line showed eye degeneration as expected ^52^ but at a lesser extent than our positive control line expressing 36 C9Orf72 expanded repeats ^53,54^ (*GMR>C9-36R*). Neither the negative control *GMR>w^-^* or the double transgenic line *GMR>NF242;TDP-43^wt^* demonstrated evidence of eye degeneration (Fig. 3h). *GMR>RGNEF;TDP-43^wt^* showed an eye phenotype different from *GMR>GFP;TDP-43^wt^* or *GMR>C9-36R* flies, that was also observed in *GMR>RGNEF* flies (Supplementary Fig. 7a) which suggests that this is an effect caused by RGNEF overexpression and is not related with TDP-43^wt^ toxicity.

Next, we studied the localization of TDP-43^wt^ and NF242 in the central brain and optical lobes (Supplementary Fig. 7b) of fixed *elav>NF242;TDP-43^wt^* and *elav>GFP;TDP-43^wt^* flies by immunofluorescence. We observed that NF242 and TDP-43^wt^ co-aggregate in the brain of flies in multiple foci in neurons (Fig. 3i-j). As expected, *elav>GFP;TDP-43^wt^* control flies showed TDP-43 foci in neurons (Supplementary Fig. 7c-d).

These results confirm that RGNEF acts as a survival factor *in vivo* and show that RGNEF and NF242 suppress the toxic motor phenotype induced by TDP-43^wt^ in flies. Also, it suggests that the co-aggregation between NF242 and TDP-43 is critical for abolishing the toxicity generated by TDP-43 overexpression in neurons.

### Ectopic NF242 expression in TDP-43 mice

The results using flies suggested a therapeutic potential for NF242. Given this, we studied the effect of the ectopic expression of NF242 in neurons using intracerebroventricular (ICV) injections of an adeno-associated virus (AAV - serotype 9) in a severe murine model of ALS (rNLS8) that expresses human TDP-43^ΔNLS^ under the regulation of a Tet-Off system^23^. AAV9 expressing GFP was used as a control.

rNLS8 mice expressing NF242 showed a significantly longer lifespan compared with the GFP-expressing animals (NF242 average: 70.28 ± 6.12 days; GFP average: 47.92 ± 6.17 days; p=0.0195; Fig. 4a). We also observed that mice injected with AAV9/NF242 had a significant improvement in clasping occurrence (NF242 average: 5.83 ± 0.27 weeks; GFP average: 4.08 ± 0.41 weeks; p=0.0075; Fig. 4b-c). As the animals injected with AAV9/NF242 were visibly more active and healthier (less kyphosis and tremor, more hydrated) compared to the mice injected with AAV9/GFP (Supplemental videos 1 - 4) in the first 5-6 weeks after doxycycline (Dox) was removed from the drinking water, we next quantified locomotor activity using an open field test^55^. Mice injected with AAV9/NF242 showed an improvement in most of the parameters evaluated for this test including total distance traveled (Fig. 4d), horizontal activity (Fig. 4e), movement time (Fig. 4f), and resting time (Fig. 4g). Vertical activity was not different between the groups (Supplementary Fig. 8a). To assess motor function and coordination, we used gait analysis (catwalk)^56,57^. We observed that mice injected with AAV9/NF242 had an improved gait phenotype (Fig. 4h) and better results in the maximum area for fore and hindlimbs with values closer to the wild type (wt) controls and significantly different from the AAV9/GFP injected rNLS8 mice (Fig. 4i-j). Additionally, swing speed for hindlimbs was significantly different compared to the AAV9/GFP injected mice controls (Fig. 4k). However, the assessment against wt controls showed a different pattern of swing speed in the transgenic mice, which is consistent with reports showing alterations in this parameter in neurodegenerative mice models with altered locomotion^57^.

**Fig. 4.**
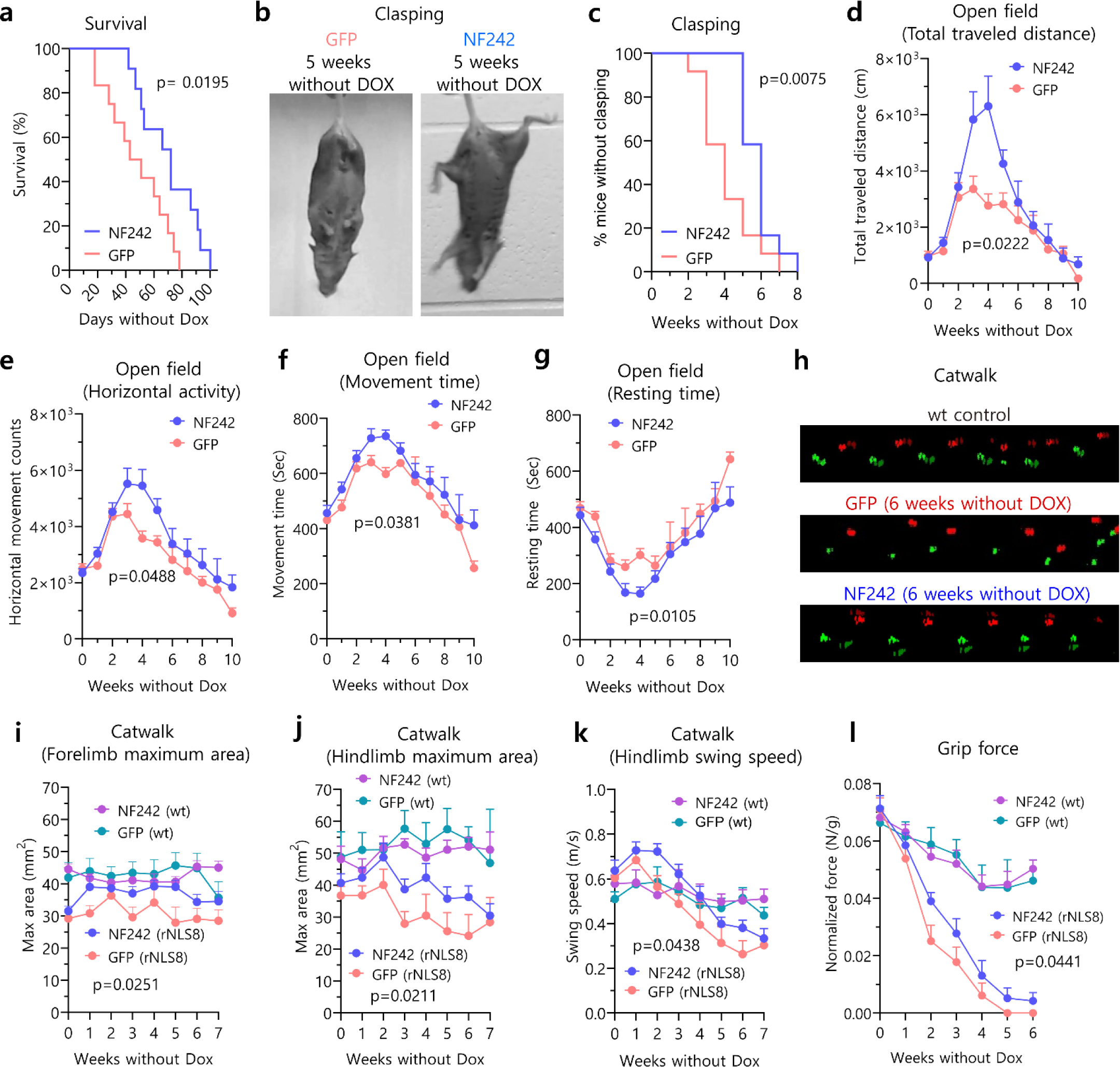
Ectopic expression of NF242 in rNLS8 mice. a, Kaplan-Meier graph showing the increased lifespan after Dox retrieval of rNLS8 mice injected with AAV9/GFP (n=12) compared to mice injected with AAV9/NF242 (n=11) (p=0.0195). b, Representative pictures showing a rNLS8 mouse injected with AAV9/GFP with clasping and a rNLS8 mouse injected with AAV9/NF242 after 5 weeks without Dox. c, Kaplan-Meier graph showing clasping quantification of rNLS8 mice injected with AAV9/GFP (n=12) or AAV9/NF242 (n=12). AAV9/NF242 injected rNLS8 mice show a significant delay in clasping occurrence (p=0.0075). d-g, Open field test comparing rNLS8 mice injected with AAV9/GFP (n=12) or AAV9/NF242 (n=12). AAV9/NF242 injected rNLS8 mice show an increase in total traveled distance (p=0.0222) (d), horizontal activity (p=0.0488) (e), movement time (p=0.0105) (f) and a decrease in resting time (p=0.0105) (g). h, Representative visualizations of gait assessment (Catwalk) that compares the improved gait pattern of an AAV9/NF242 injected rNLS8 mouse with an AAV9/GFP injected rNLS8 mouse at 6 weeks without Dox. Wild-type mouse shows normal gait. i-k, Catwalk quantification comparing rNLS8 and wt mice injected with AAV9/GFP or AAV9/NF242 (n=12 for each group of rNLS8 mice; n=6 for each group of wt mice). The AAV9/NF242 injected rNLS8 mice show an improvement in the forelimb maximum area (p=0.0251) (i), the hindlimb maximum area (p=0.0211) (j), and the hindlimb swing speed (p=0.0438) (k). l, Grip force experiment showing that the AAV9/NF242 injected rNLS8 mice (n=12) have a slight increase in the force compared to the AAV9/GFP injected mice (n=12) (p=0.0441). Wt mice injected with AAV9/GFP (n=6) or AAV9/NF242 (n=6) are shown as healthy grip force controls.

We did not observe a difference for fore and hindlimbs stride length (Supplementary Fig. 8b-c) or forelimb swing speed (Supplementary Fig. 8d). When we analyzed the strength of the mice using a grip force test, we also observed a better performance for mice injected with AAV9/NF242 (Fig. 4l). We did not observe a difference in the balance using rotarod test^58^ (Supplementary Fig. 8e) or in the weight of the mice injected with AAV9/NF242 compared to AAV9/GFP (Supplementary Fig. 8f).

Fluorescence staining showed that the AAV9/GFP and AAV9/NF242 were efficiently transduced in the brain of wild-type mice (Supplementary Fig. 9a-b). The same was observed when the viruses were injected in rNLS8 mice (Supplementary Fig. 9c-d). When we analyzed the spinal cord of rNLS8 mice injected with AAV9/NF242, we observed a high efficiency of transduction (Supplementary Fig. 9e). Neurons expressing NF242 in the spinal cord and brain cortex of rNLS8 mice observed at high magnification with confocal microscopy or using super-resolution stimulated emission depletion (STED) microscopy showed extensive co-localization and co-aggregation with TDP-43^ΔNLS^ (Fig. 5a-b). When we analyzed the expression of the neuroinflammatory markers in spinal cord, we observed a significant reduction in the levels of the astrogliosis marker GFAP (Fig. 5c; reduction of 59.9%; p=0.0033; Fig 5e) and the microgliosis marker Iba1 (Fig. 5d; reduction of 74.4%; p=0.0341; Fig. 5f) in the NF242 expressing mice.

**Fig. 5.**
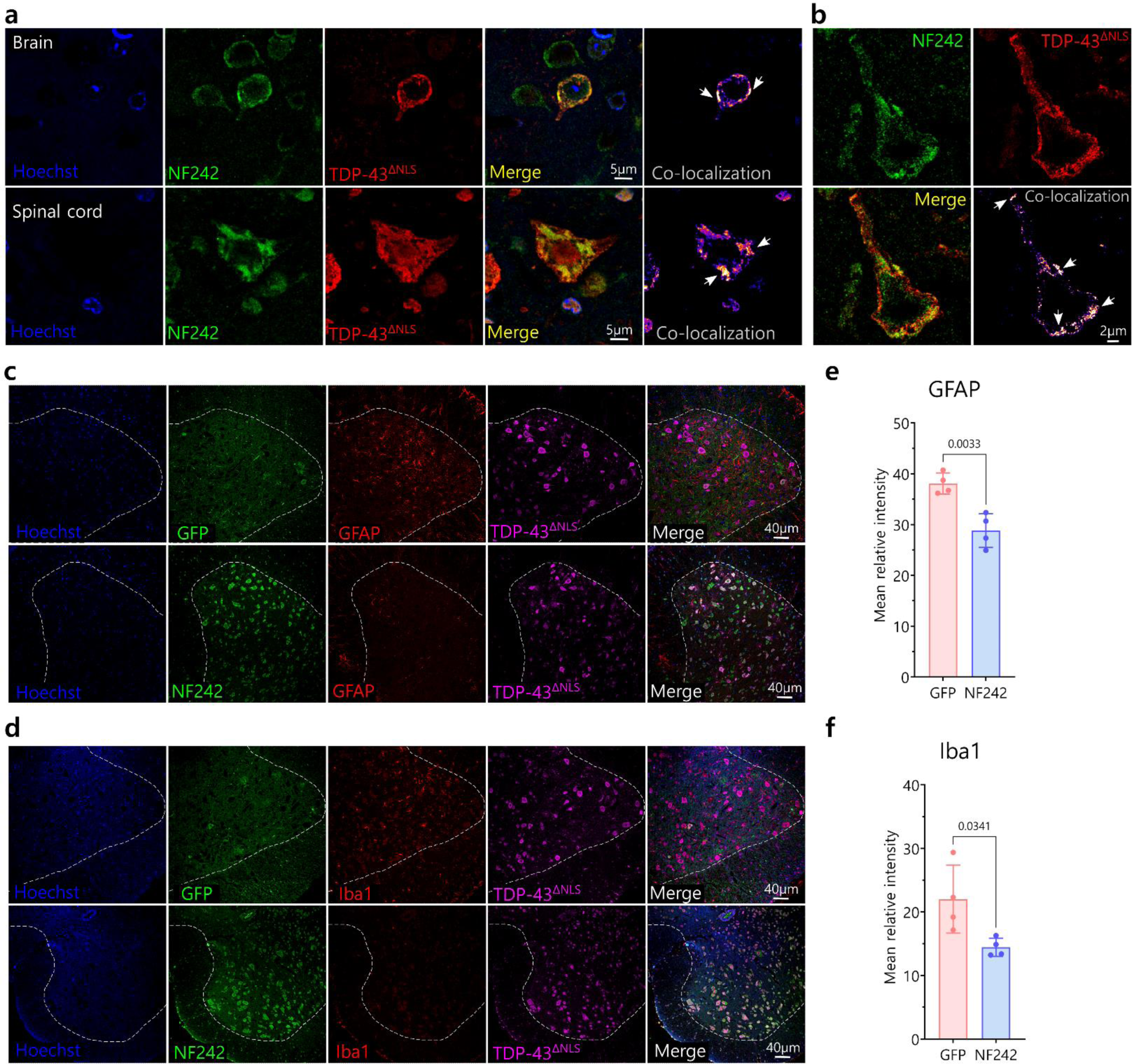
Pathology of rNLS8 mice expressing ectopic NF242 at week 3. a, High magnification confocal images showing the co-localization and co-aggregation (indicated by white arrows) between NF242 and TDP-43^ΔNLS^ in the brain cortex (cortical layer II-III) and spinal cord of a rNLS8 mouse injected with AAV9/NF242 after 3 weeks without Dox. b, Super-resolution STED microscopy images showing in detail the co-aggregation (indicated by white arrows) between NF242 and TDP-43^ΔNLS^ in the brain cortex (cortical layer II-III) of a rNLS8 mouse injected with AAV9/NF242 after 3 weeks without Dox c-d, Representative immunofluorescences of rNLS8 mice injected with AAV9/GFP and AAV9/NF242 showing the decrease in the amount of GFAP (c) and Iba1 (d) in the spinal cord. The anterior gray horn is separated from the white matter by a dashed white line. e-f, Quantification showing the reduction of the levels of GFAP (e, p=0.0033) and Iba1 (f, p=0.0341) in the ventral horns of the lumbar spinal cord of rNLS8 mice injected with AAV9/GFP and AAV9/NF242, after 3 weeks without Dox (n=4).

These results demonstrate that the ectopic expression of NF242 in neurons of mice brain and spinal cord using AAVs improves the lifespan and motor phenotype of a severe and fast-deteriorating model of ALS based on TDP-43 dysregulation.

## Discussion

Here, we show that the pathological phenotype induced by TDP-43 overexpression is suppressed in flies, and ameliorated in mice by an amino-terminal fragment of the RNA-binding protein RGNEF/p190RhoGEF (NF242). This is the first report of a protein fragment that naturally binds TDP-43 and has a therapeutic effect that includes improvement of motor phenotype, increased lifespan, and reduction of neuroinflammatory markers in a murine ALS model.

Our results support that the interaction and specific co-aggregation between NF242 and TDP-43 are key to the protective effect of NF242 against the toxicity induced by TDP-43^wt^ and TDP-43^ΔNLS^ in two *in vivo* models. Since NF242 competes with RNA for the binding site of TDP-43, NF242 might be blocking the gain of toxic function of TDP-43 generated by sequestering RNA into the aggregates^59^. The moderate effect of NF242 over TDP-43 activity and the robust effect in pathological models of TDP-43 overexpression *in vivo* without deleterious effect in controls expressing NF242 alone, suggests that NF242 has a higher affinity for pathological TDP-43 than for normal TDP-43, leading to co-aggregation.

Thus far, the RRMs of TDP-43 have been targeted for potential therapeutic approaches in a few *in vitro* and *in vivo* studies using flies and mice. This includes the utilization of small molecules such as compounds containing extended planar aromatic moieties^60^, ATP^61^, the chemical rTRD01^62^, and an antibody against the RRM1 of TDP-43^63^. These data support our findings demonstrating that targeting the RRMs domains of TDP-43 can improve the phenotype of TDP-43 proteinopathies.

In the rNLS8 mice, a well-studied ALS animal model^23,64–66^, the mitigation effect of the AAV9/NF242 over the motor phenotype lasted 5-6 weeks on average. The progression of signs thereafter could be explained by two reasons. First, the high and permanent expression of TDP-43^ΔNLS^ in the model and the spreading of the TDP-43 pathology beyond the cells expressing NF242. Specifically, we observed a significant increase of TDP-43 pathology in the cortical layer I at 6 weeks without Dox in rNLS8 mice injected with AAV9/NF242. This increase of TDP-43 pathology was correlated with an increase of GFAP in the cortical layers I and II-III and an extensive co-localization of TDP-43^ΔNLS^ with GFAP mainly in the cortical layer I (Fig. 6), suggesting the spreading of TDP-43 pathology to astrocytes after several weeks of TDP-43^ΔNLS^ neuronal expression. Second, the suppression of the endogenous murine TDP-43 expression in the rNLS8 mice^23^; the loss of function is an inherent pathological factor in the model that cannot be accounted for by our therapeutic approach.

**Fig. 6.**
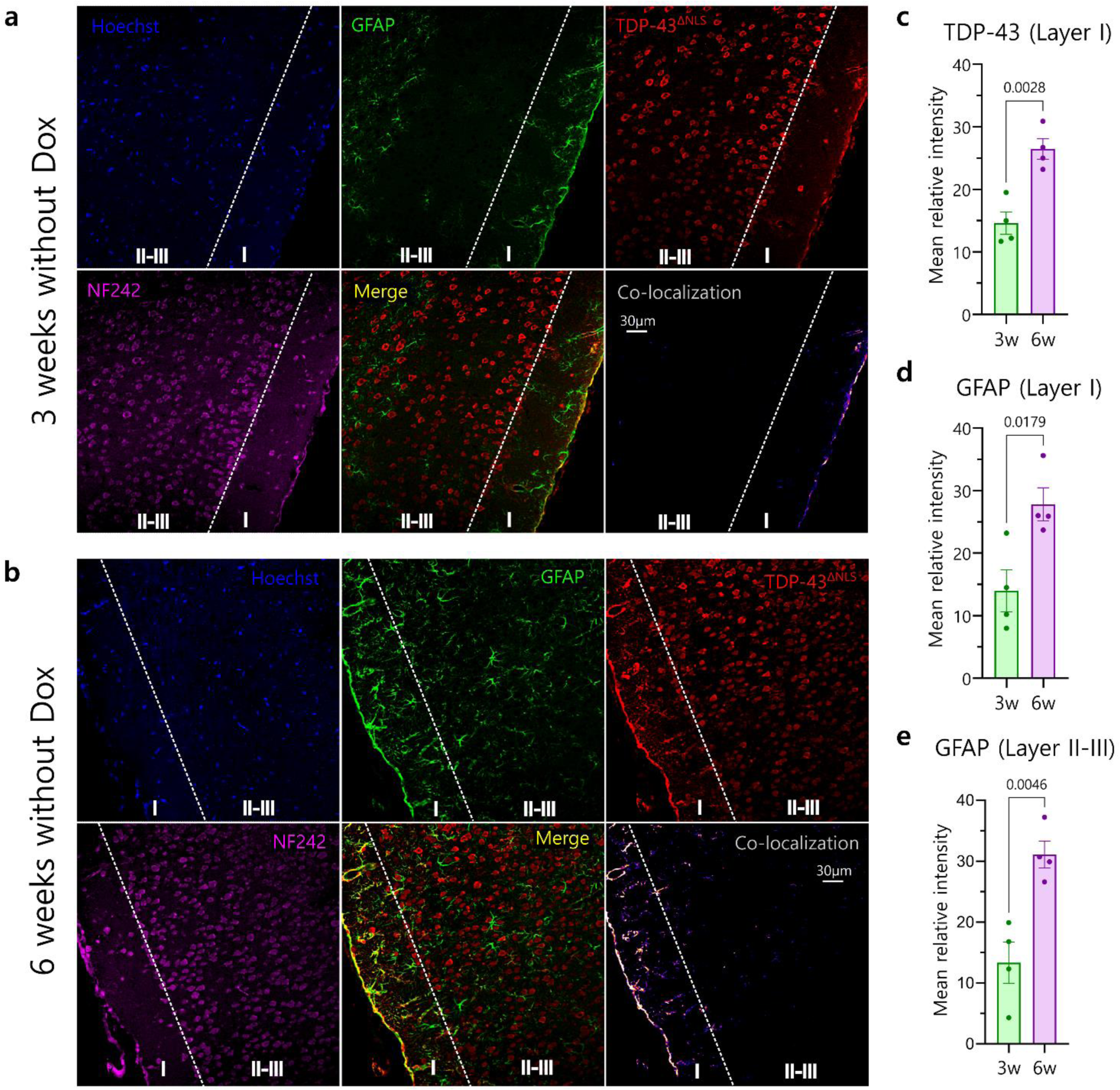
Comparison of rNLS8 mice brain expressing ectopic NF242 between week 3 and 6. a, Localization analysis of GFAP and TDP-43ΔNLS in the cortex of rNLS8 mouse injected with AAV9/NF242, after 3 weeks without Dox. Notice the low levels of TDP-43^ΔNLS^ in the cortical layer I of the brain and the absence of co-localization between TDP-43ΔNLS and GFAP in that region. b, Localization analysis of GFAP and TDP-43^ΔNLS^ in the cortex of rNLS8 mouse injected with AAV9/NF242, after 6 weeks without Dox. Notice the spreading of TDP-43 pathology in the cortical layer I of the brain and the intensive co-localization between GFAP and TDP-43^ΔNLS^ in that region. c, Quantification of the increase of TDP-43^ΔNLS^ amount in the cortical layer I of the brain from 3 to 6 weeks without Dox (n=4, p=0.0028). d, Quantification of the increase of GFAP amount in the cortical layer I of the brain from 3 to 6 weeks without Dox (n=4, p=0.0179). e, Quantification of the increase of GFAP amount in the cortical layer II-III of the brain from 3 to 6 weeks without Dox (n=4, p=0.0046). The cortical layer I and II-III are indicated in all images.

Considering that the rNLS8 mice have previously shown resistance to other therapeutic approaches, such as riluzole treatment^24^, MMP-9 reduction^67^, or miR-23a suppression^68^, the more encouraging evidence that we show in this work highlights the therapeutic potential of our approach after modifying the phenotype of this severe murine model of ALS.

In conclusion, our study suggests that an analogous therapeutic strategy expressing NF242 or a biologically active fragment of NF242 could be promising in humans affected by TDP-43 proteinopathies. The fact that this approach uses a fragment of a protein already expressed in humans, suggests that the secondary effects associated with the use of therapeutic antibodies^69,70^ could be minimized or avoided.

A potential treatment using this TDP-43’s gain-of-function targeting approach might need to be combined with drugs that target its loss-of-function, and potentially with drugs focused on different targets, such as autophagy^71^. Currently, it seems we are on the edge of a new era for the developing of treatments for neurodegenerative diseases such as ALS and frontotemporal dementia.

## Acknowledgements

The authors are grateful for the ongoing collaborations with Drs. Robert Bowser and Janice Robertson as members of an RGNEF study group. Stocks obtained from the Bloomington Drosophila Stock Center (NIH P40OD018537) were used in this study.

## Author contributions

C.A.D. conceived the project, performed and analyzed *in vitro* experiments, performed and analyzed *in vivo* experiments with flies and analyzed experiments with mice, analyzed and checked the overall quality of the data, prepared the illustrations, and drafted the manuscript; D.C.M. performed and analyzed *in vitro* and pathology experiments, collaborated with the overall analysis of data, and drafted the manuscript; V.N. performed *in vitro* experiments, performed and analyzed the *in vivo* experiments with mice and contributed with the draft; C.M. prepared, performed, and analyzed the SPR experiments and contributed with the draft; R.S. performed the protein-protein interaction modeling analysis and contributed with the draft; T.A.L. performed *in vivo* experiments with flies; H.A. contributed to *in vitro* experiments; B.W. contributed to *in vivo* experiments with flies; K.S.S. purified full TDP-43 for the SPR experiments; A.S., E.B., M.J., and J.K. contributed technically, with insight, and reviewed the final version of the paper; M.J.S. secured funding, oversaw the project development, reviewed and approved the final submission of the manuscript.

## Additional information

Supplementary information is available for this paper. Correspondence and request for materials should be addressed to Dr. Cristian Droppelmann.

