## Supplementary Material for "Mitigation of TDP-43-induced toxic phenotype by expression of RGNEF N-terminal fragment in ALS models"

#### Supplementary Figure 1

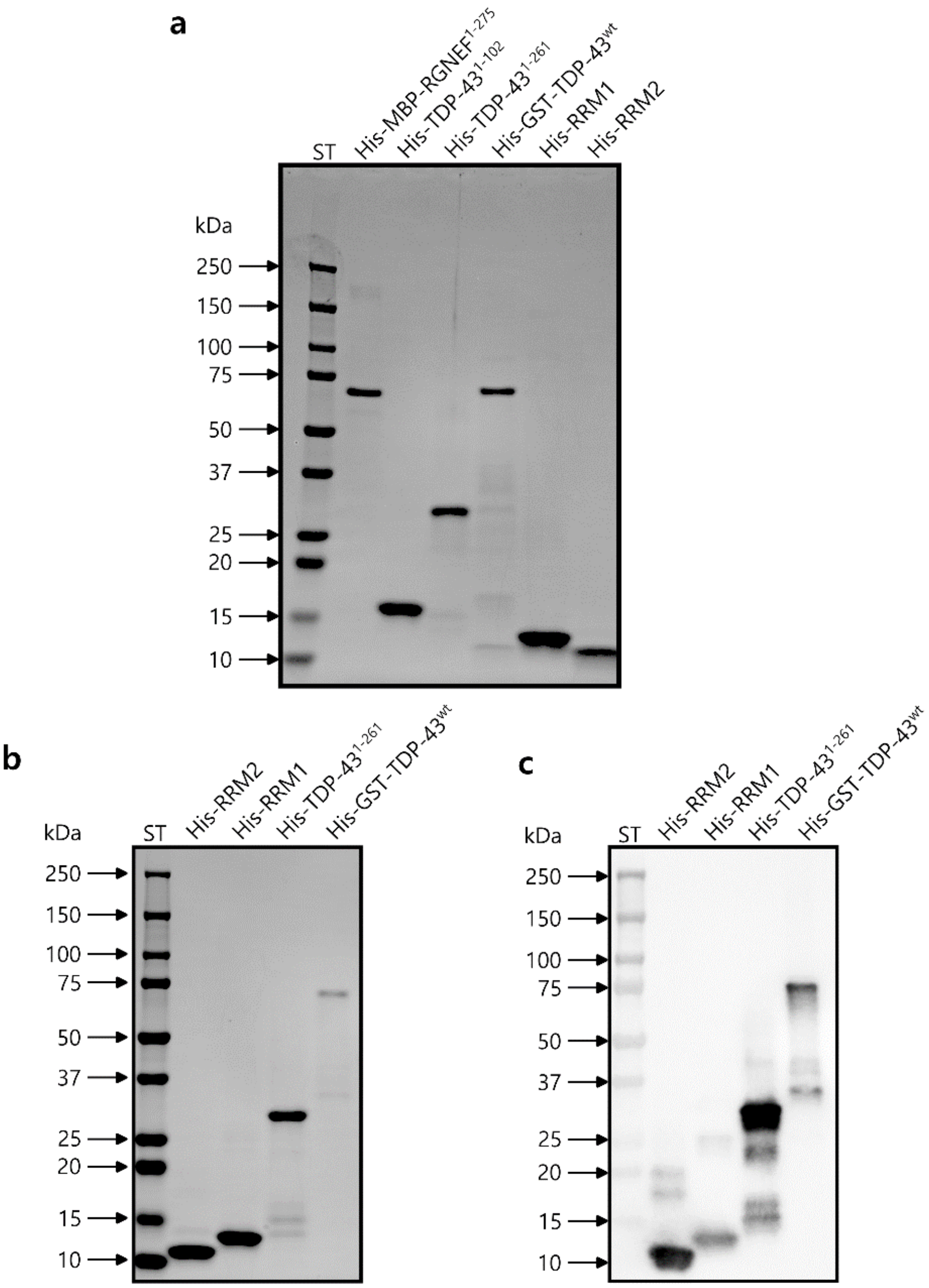

**Supplementary Fig. 1 | Gel electrophoresis of recombinant proteins purified for SPR.** **a**, approximately 1 µg of purified His-MBP-RGNEF$^{1\text{-}275}$, His-TDP-43$^{1\text{-}102}$, His-TDP-43$^{1\text{-}261}$, His-GST-TDP-43$^{wt}$, His-RRM-1, and His-RRM-2 were loaded. **b**, approximately 0.6 µg of purified His-TDP-43$^{1\text{-}261}$, His-RRM-1, and His-RRM-2 samples, and approximately 0.14 µg of purified His-GST-TDP-43$^{wt}$ sample was loaded. **c**, The same samples (40% less protein) from (c) were also loaded onto a separate precast gradient gel and then transferred to nitrocellulose membrane for western blot analysis using a TDP-43 antibody. The molecular weight standard (ST) was imaged using the colorimetric channel and merged with the chemiluminescent image. His: 6x histidine tag; GST: Glutathione S-transferase.

#### Supplementary Figure 2

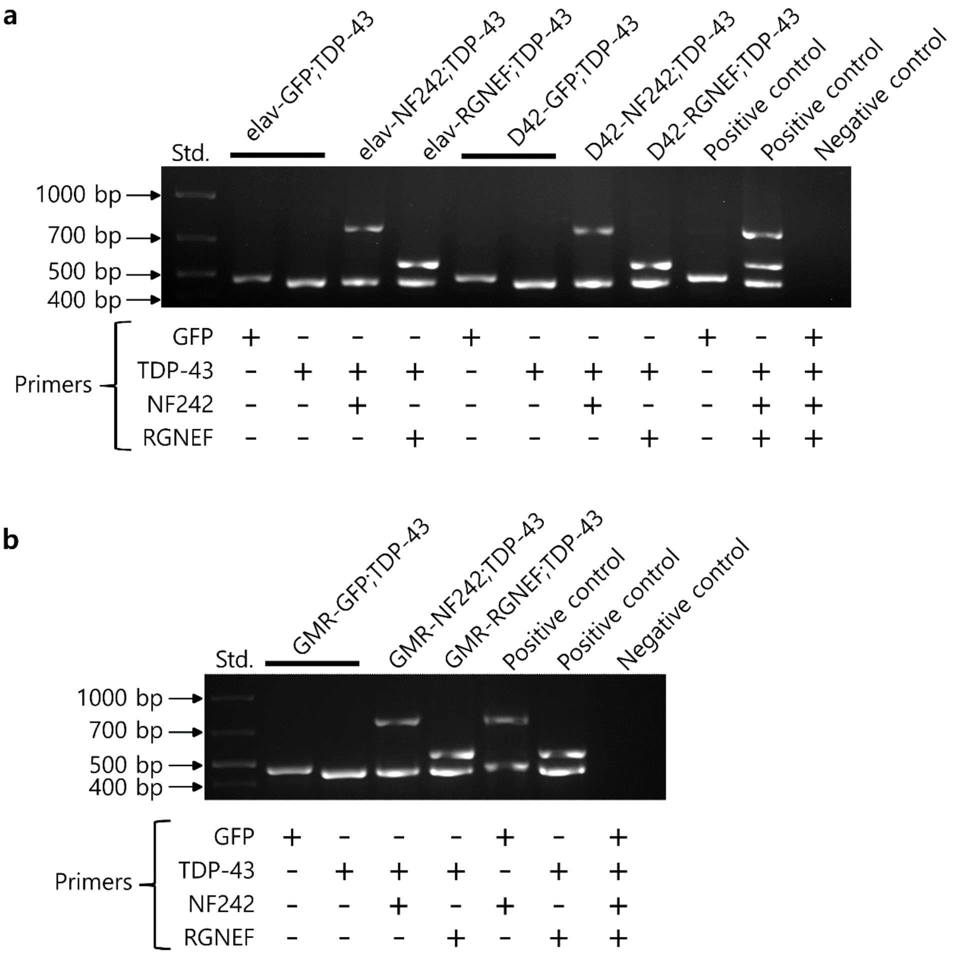


**Supplementary Fig. 2 | Expression of genes of interests in transgenic drosophila. a**, Expression of GFP, TDP-43, NF242 and RGNEF in double transgenic flies under the elav and D42 drivers. **b**, Expression of GFP, TDP-43, NF242 and RGNEF in double transgenic flies under the GMR driver. As positive controls were used plasmids containing the genes of interest.

### Supplementary Figure 3

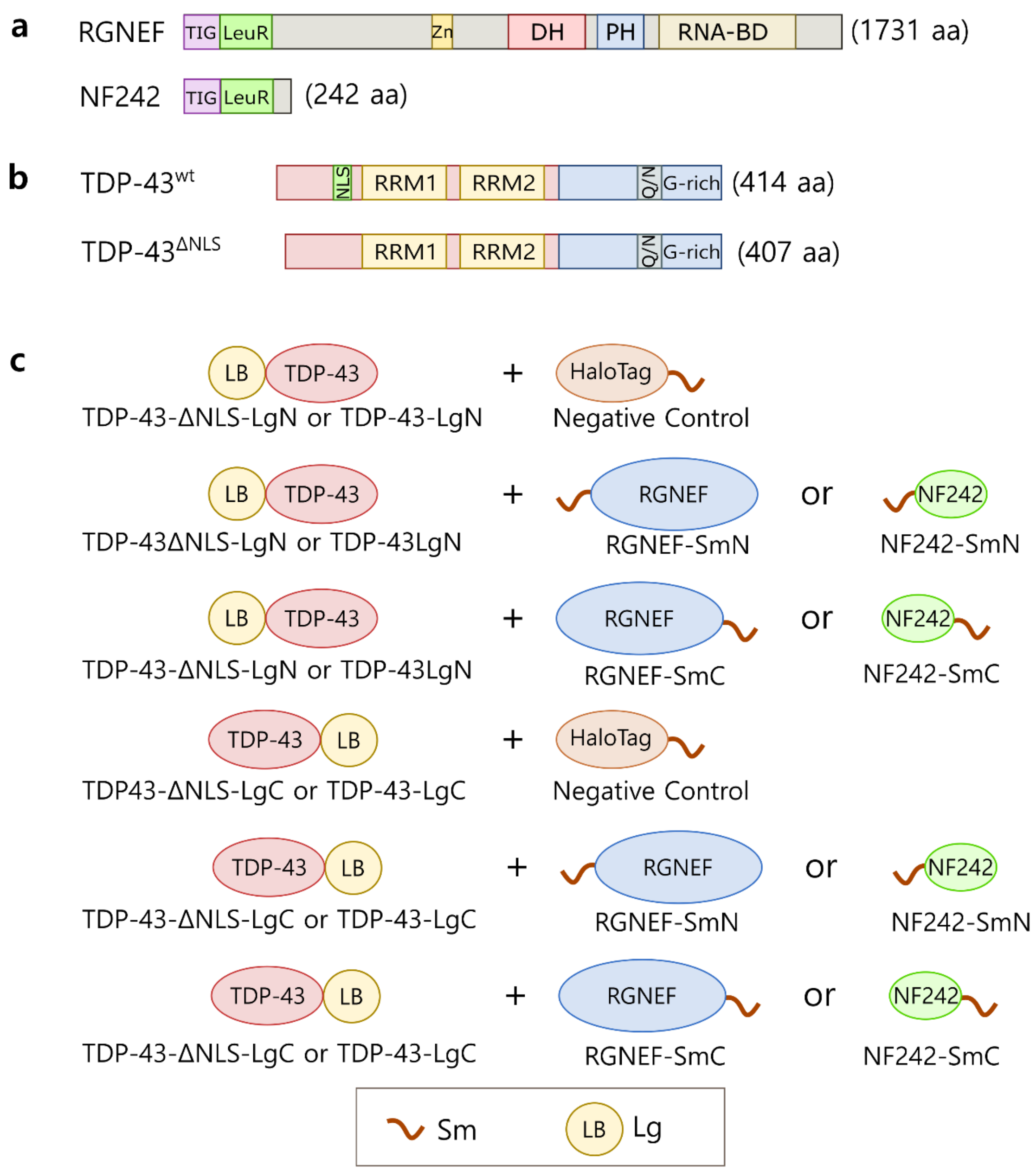

**Supplementary Fig. 3 | Constructs used for NanoBiT experiments.** **a**, Schematics of RGNEF and NF242 sequences used in this study. TIG: IPT/TIG domain; LeuR: Leucine-rich domain; Zn: cysteine-rich Zinc binding domain; DH: Dbl homology domain; PH: Pleckstrin homology domain; RNA-BD: RNA-binding domain. **b**, Schematic of the TDP-43$^{wt}$ and TDP-43$^{\Delta NLS}$ sequences used in this study. NLS: Nuclear localization signal; RRM1: RNA recognition motif 1; RRM2: RNA recognition motif 2; Q/N: Glutamine/Asparagine-rich segment; G-rich: Glycine rich region. **c**, Schematic of the NanoBiT constructs used in this study. Sm: Small subunit of luciferase (11 amino acids); Lg: Large subunit of luciferase (17.6 kDa); N: Amino-terminal; C: Carboxy-terminal.

### Supplementary Figure 4

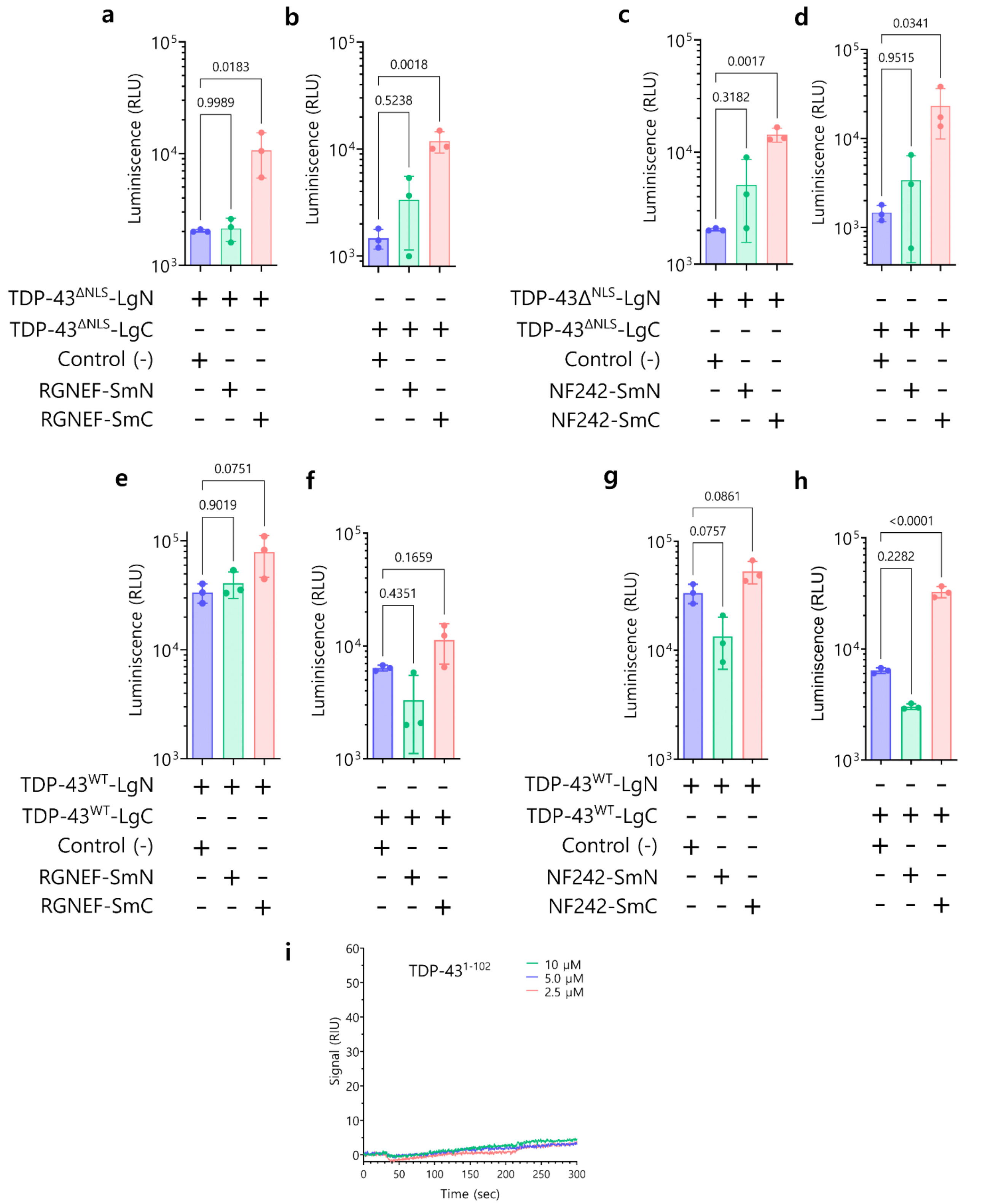

**Supplementary Fig. 4 | Interaction between RGNEF or NF242 with TDP-43 using complementation reporter assay (NanoBiT). a**, Interaction was observed between TDP-$43^{\Delta NLS}$ fused with the large subunit of luciferase (Lg) in the amino-terminal end (TDP-$43^{\Delta NLS}$-LgN) and RGNEF fused with small subunit of luciferase (Sm) in the carboxy-terminal end (RGNEF-SmC; p=0.0183) but not in the amino-terminal end (RGNEF-SmN; p=0.9989). **b**, Interaction was observed between TDP-$43^{\Delta NLS}$ fused with Lg in the carboxy-terminal end (TDP-$43^{\Delta NLS}$-LgC) and RGNEF fused with Sm in the carboxy-terminal end (RGNEF-SmC; p=0.0018) but not in the amino-terminal end (RGNEF-SmN; p=0.5238). **c**, Interaction was observed between TDP-$43^{\Delta NLS}$ fused with Lg in the amino-terminal end (TDP-$43^{\Delta NLS}$-LgN) and NF242 fused with Sm in the carboxy-terminal end (NF242-SmC; p=0.0017) but not in the amino-terminal end (NF242-SmN; p=0.3182). **d**, Interaction was observed between TDP-$43^{\Delta NLS}$ fused with Lg in the carboxy-terminal end (TDP-$43^{\Delta NLS}$-LgC) and NF242 fused with Sm in the carboxy-terminal end (NF242-SmC; p=0.0341) but not in the amino-terminal end (NF242-SmN; p=0.9515). **e**, No interaction was observed between TDP-$43^{wt}$ fused with Lg in the amino-terminal end (TDP-$43^{wt}$-LgN) and RGNEF fused with Sm in the carboxy-terminal end (RGNEF-SmC; p=0.0751) or in the amino-terminal end (RGNEF-SmN; p=0.9019). **f**, No interaction was observed between TDP-$43^{wt}$ fused with Lg in the carboxy-terminal end (TDP-$43^{wt}$-:LgC) and RGNEF fused with Sm in the carboxy-terminal end (RGNEF-SmC; p=0.1659) or in the amino-terminal end (RGNEF-SmN; p=0.4351). **g**, No interaction was observed between TDP-$43^{wt}$ fused with Lg in the amino-terminal end (TDP-$43^{wt}$-LgN) and NF242 fused with Sm in the carboxy-terminal end (NF242-SmC; p=0.0861) or in the amino-terminal end (NF242-SmN; p=0.0757). **h**, Interaction was observed between TDP-$43^{wt}$ fused with Lg in the carboxy-terminal end (TDP-$43^{wt}$-LgC) and NF242 fused with Sm in the carboxy-terminal end (NF242-SmC; p<0.0001) but not in the amino-terminal end (NF242-SmN; p=0.2282). N=3 and 6 technical replicates for each experiment. These results demonstrated that NF242 and TDP-43 fused in the carboxy-terminal end to the luciferase fragments are the best pair for interaction experiments. This configuration was used for the experiments presented in Fig.1. **i**, Representative SPR sensorgrams demonstrating the absence of interaction between His-TDP-$43^{1-102}$ (analyte) and His-MBP-$RGNEF^{1-275}$ (ligand) at different concentrations of His-TDP-$43^{1-102}$ (n=3).

#### Supplementary Figure 5

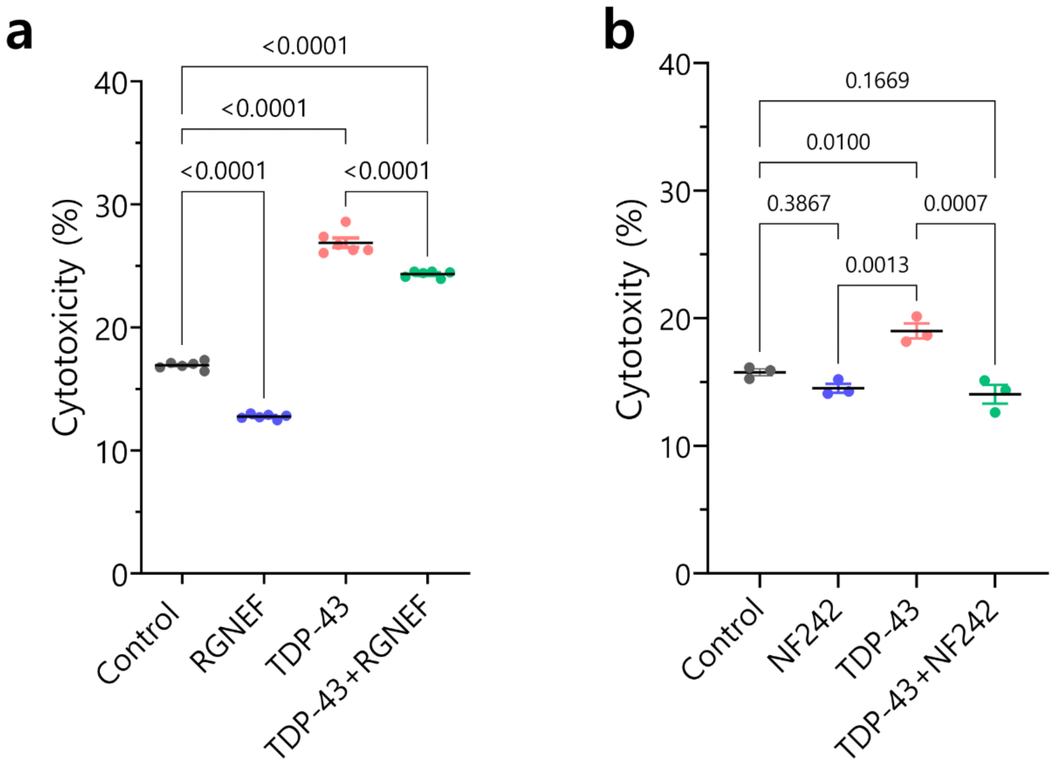


**Supplementary Fig. 5 | Cytotoxicity assay of cells transfected with RGNEF, NF242 and TDP-43. a-b**, The increase of the cytotoxicity induced by TDP-43 overexpression is reduced with the co-expression of RGNEF (n=6) (**a**) or NF242 (n=3) (**b**). Controls cells transfected with empty vector (control) or vector expressing RGNEF or NF242 alone. RGNEF alone reduced the cytotoxicity in consistence with our previous reports.

#### Supplementary Figure 6

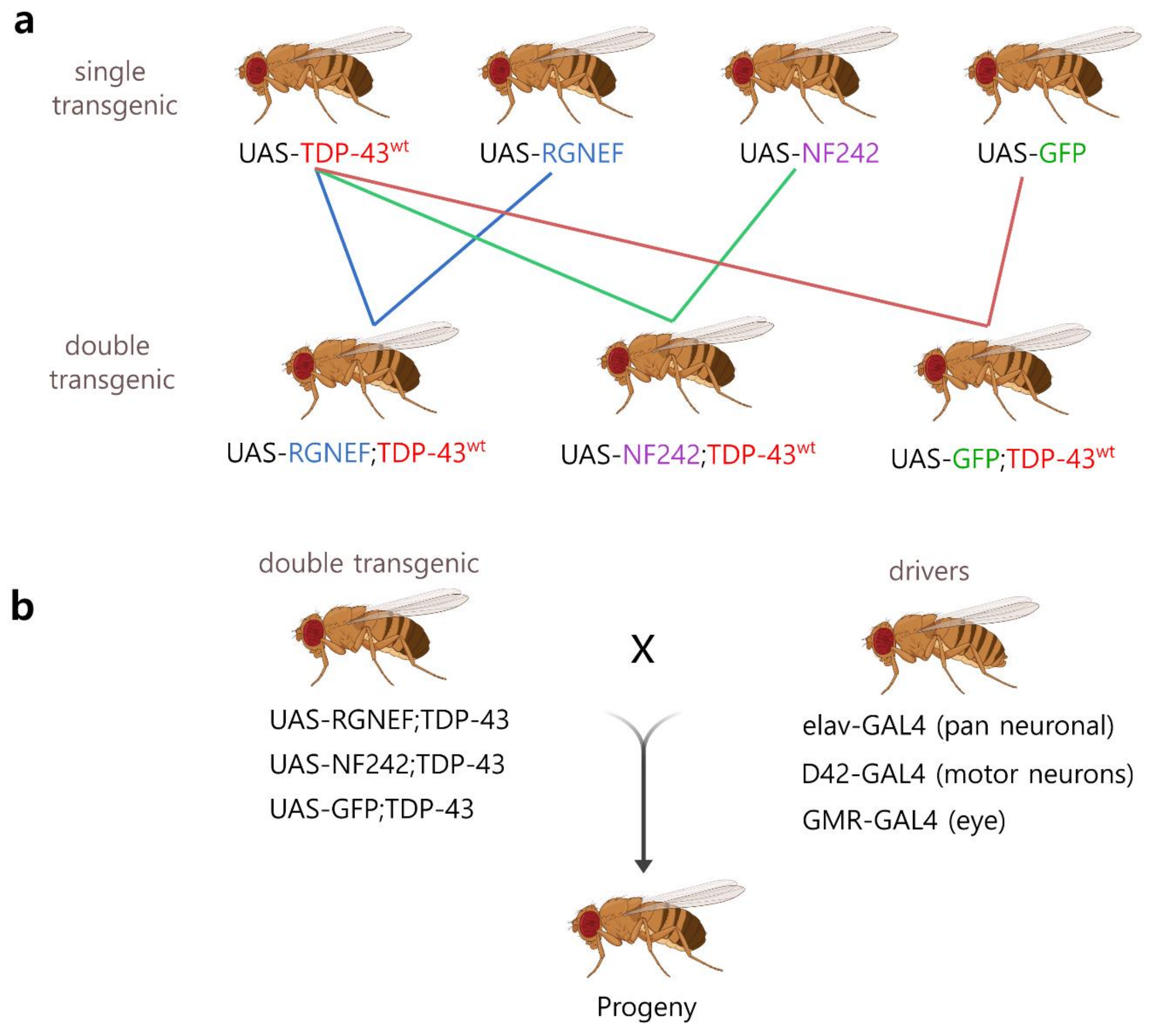


**Supplementary Fig. 6 | Transgenic flies used in this study.** **a**, *UAS-TDP-43*, *UAS-RGNEF*, and *UAS-NF242* single transgenic flies were generated for this study. The *UAS-GFP* fly was obtained from Bloomington Drosophila Stock Center. Flies were crossed to generate double transgenic effector lines: *UAS-RGNEF;TDP-43$^{wt}$*, *UAS-NF242;TDP-43$^{wt}$*, and *UAS-GFP;TDP-43wt*. **b**, Schematic showing the crosses between effector and driver lines performed in this study.

#### Supplementary Figure 7

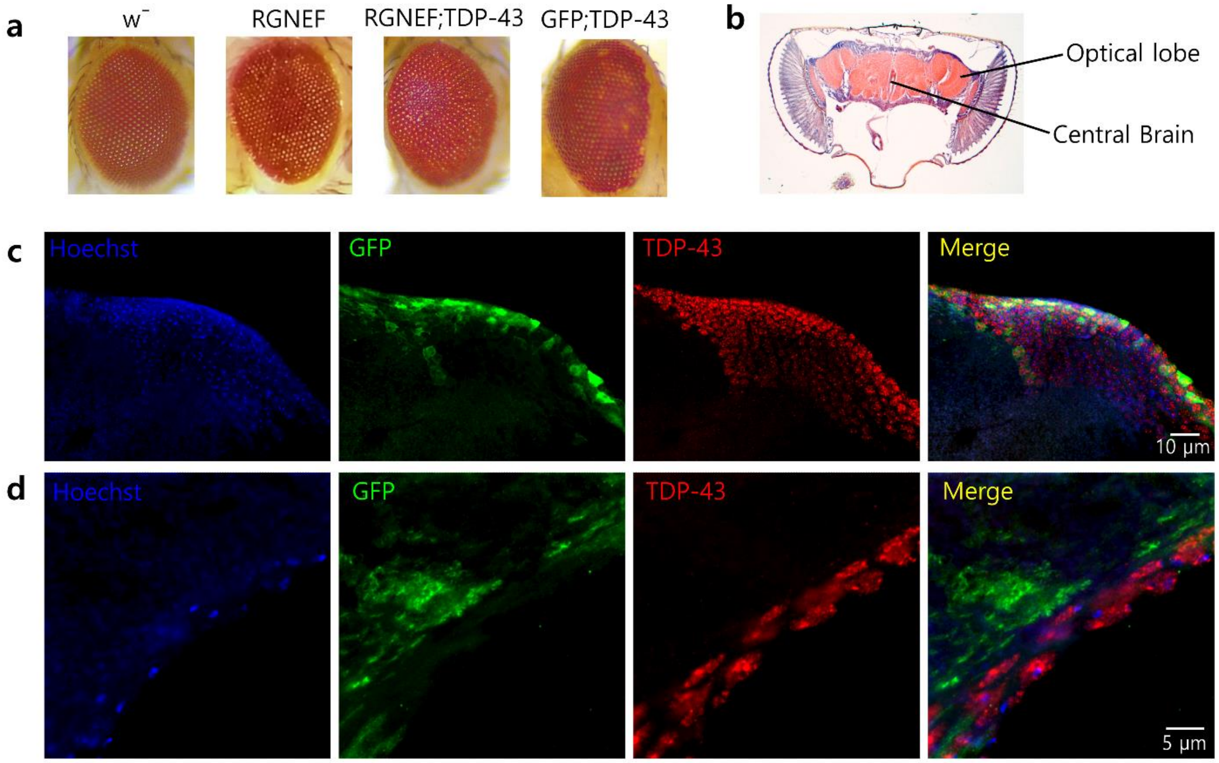


**Supplementary Fig. 7 | Co-expression of RGNEF or NF242 with TDP-43 in fruit flies (additional data). a**, Representative images showing the eye phenotype of lines *GMR>w⁻* (negative control), *GMR>RGNEF*, *GMR>RGNEF;TDP-43$^{wt}$*, and *GMR>GFP;TDP-43$^{wt}$*. Both *GMR>RGNEF*, *GMR>RGNEF;TDP-43$^{wt}$*, show a distinctive eye phenotype that is different from the pathological *GMR>GFP;TDP-43$^{wt}$*. **b**, Hematoxylin-eosin staining showing an adult *Drosophila* head. The brain regions, including the optical lobes, analyzed in this study are indicated. **c**, Immunofluorescence of adult *elav>GFP;TDP-43$^{wt}$* fly brain showing the localization GFP and TDP-43 in neurons. **d**, Higher magnification of adult *elav>GFP;TDP-43$^{wt}$* fly brain showing the aggregation of TDP-43 in neurons.

#### Supplementary Figure 8

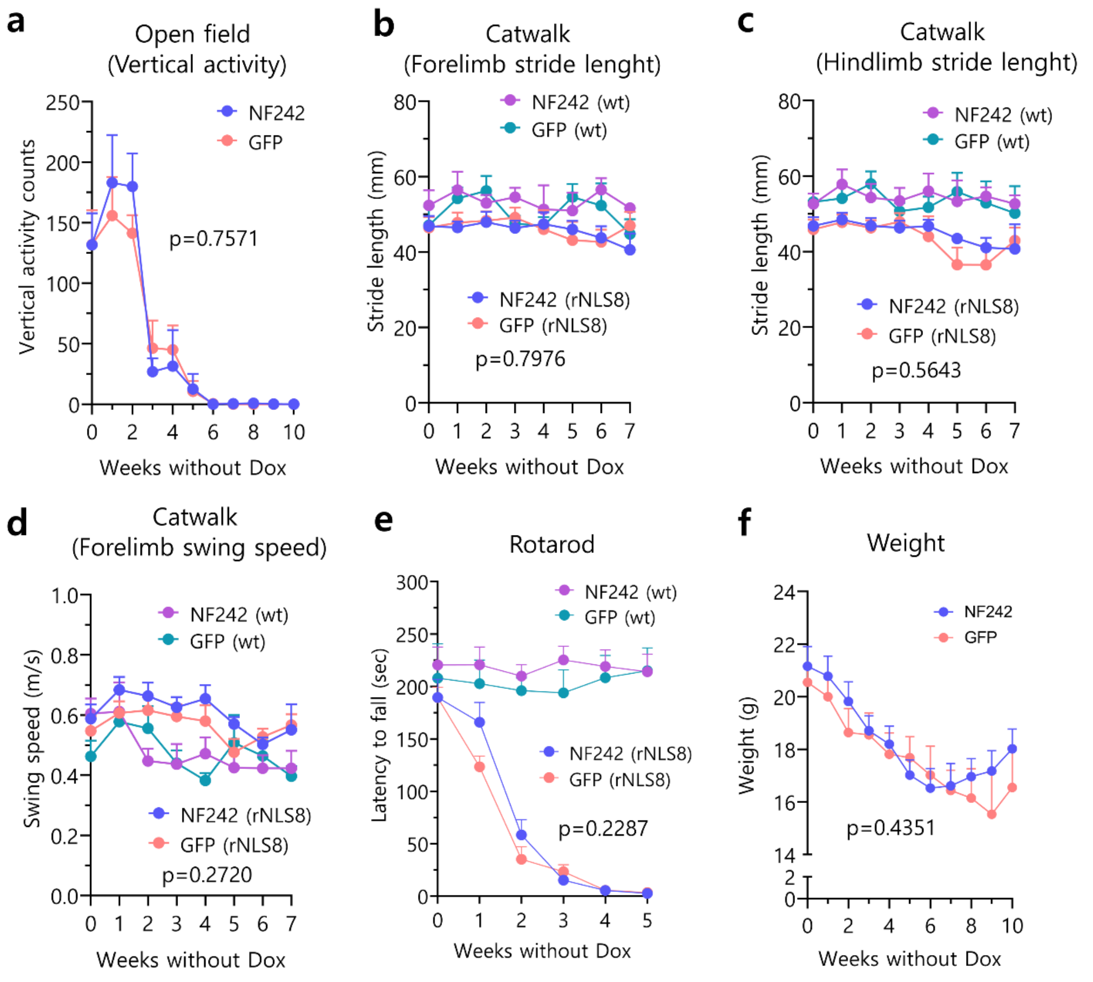


**Supplementary Fig. 8 | Ectopic expression of NF242 in rNLS8 mice (Additional data).** **a**, Open field test comparing rNLS8 mice injected with AAV9/GFP (n=12) or AAV9/NF242 (n=12) shows no difference in vertical activity between the two groups (p=0.7571). **b-d**, Catwalk quantification comparing rNLS8 and wt mice injected with AAV9/GFP or AAV9/NF242 (n=12 for each group of rNLS8 mice; n=6 for each group of wt mice). The two rNLS8 groups show no difference in forelimb stride length (p=0.7976) (**b**), hindlimb stride length (p=0.5643) (**c**), and forelimb swing speed (p=0.2720) (**d**). **e**, Rotarod test show no difference between rNLS8 mice injected with AAV9/GFP (n=12) or AAV9/NF242 (n=12) (p=0.2741). Wt mice injected with AAV9/GFP (n=6) or AAV9/NF242 (n=6) are shown as healthy controls. **f**, Weight of rNLS8 mice injected with AAV9/GFP (n=10) or AAV9/NF242 (n=9). No significant difference was found between the groups (p=0.4351).

### Supplementary Figure 9

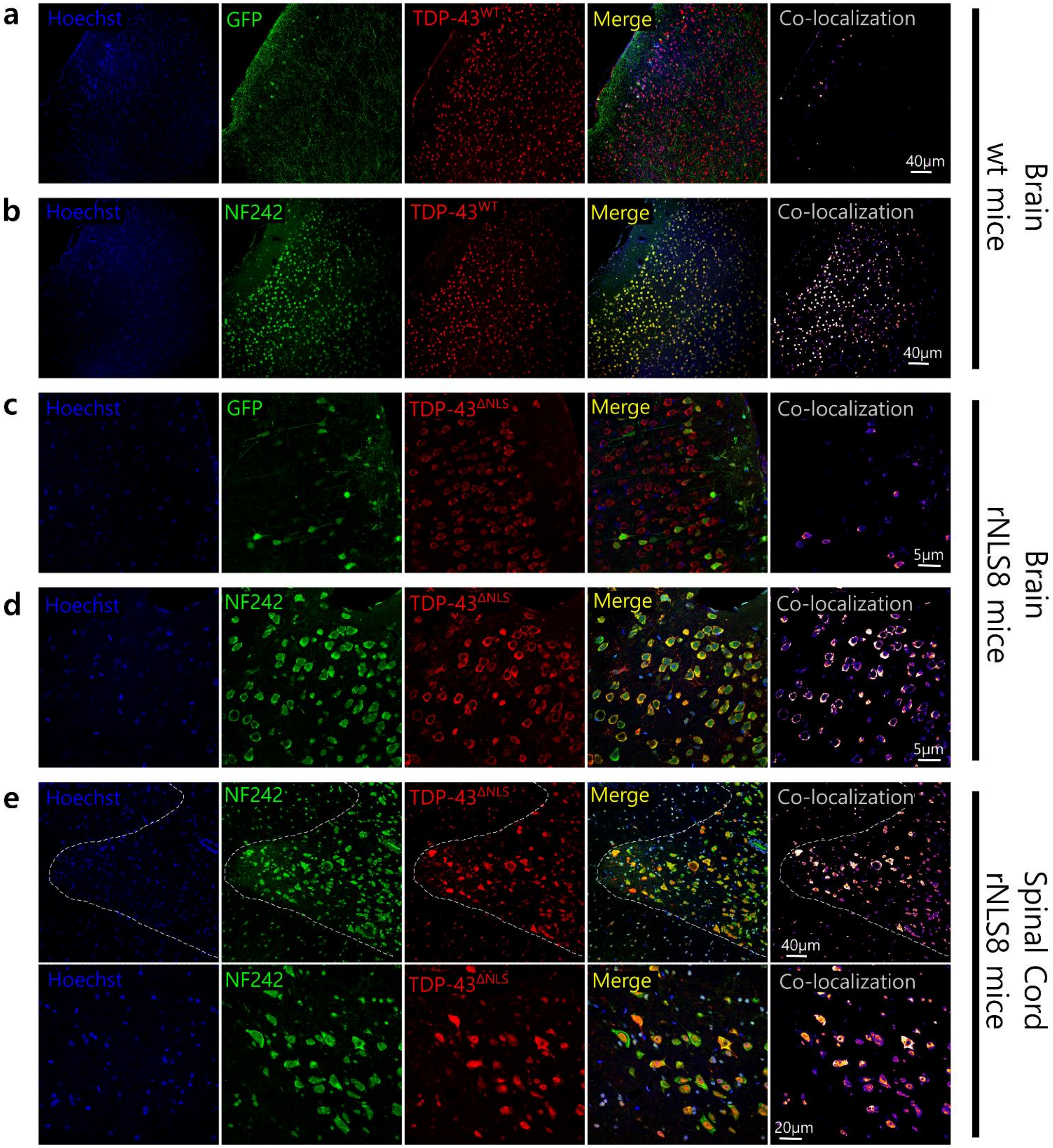

**Supplementary Fig. 9 | Pathology of rNLS8 mice expressing ectopic NF242 (Additional data). a**, Expression of GFP and endogenous TDP-$43^{wt}$ in the brain cortex (cortical layer II-III) of wild-type mouse injected with AAV9/GFP, after 3 weeks without Dox. **b**, Expression of NF242 and endogenous TDP-$43^{wt}$ in the brain cortex (cortical layer II-III) of wt mouse injected with AAV9/NF242, after 3 weeks without Dox. **c**, Expression of GFP and TDP-$43^{\Delta NLS}$ in the brain cortex (cortical layer II-III) of rNLS8 mouse injected with AAV9/GFP, after 3 weeks without Dox. **d**, Co-localization of NF242 with TDP-$43^{\Delta NLS}$ in the brain cortex (cortical layer II-III) of rNLS8 mouse injected with AAV9/NF242, after 3 weeks without Dox. **e**, Two magnifications showing the co-localization of NF242 with TDP-$43^{\Delta NLS}$ in the spinal cord of rNLS8 mouse injected with AAV9/NF242, after 3 weeks without Dox. The gray matter is separated from the white matter by a dashed white line.

**Supplementary Table 1**

| Reagent or Resource | Source | Catalog number |
|---|---|---|
| Antibodies | | |
| Anti-flag (goat) | Novus Biologicals | NB600-344 |
| Anti-GFP (rabbit) | Abcam | ab6673 |
| Anti-TDP-43 (rabbit) | Proteintech | 10782-2-AP |
| Anti-TDP-43 (mouse) | Abcam | ab104223 |
| Anti-Iba1 (rabbit) | Wako | 019-197-41 |
| Anti-GFAP (mouse) | BD | 556330 |
| Anti-rabbit HRP conjugated (swine) | Dako | P0399 |
| Anti-goat Alexa 488 (donkey) | Invitrogen | A11055 |
| Anti-rabbit Alexa 555 (donkey) | Invitrogen | A31572 |
| Anti-mouse Alexa 555 (donkey) | Invitrogen | A31570 |
| Anti-rabbit Alexa 594 (donkey) | Invitrogen | A21207 |
| Anti-rabbit Alexa 680 (donkey) | Invitrogen | A10043 |
| Chemicals and kits | | |
| Hoechst 32258 | Sigma | B-1155 |
| DAPI | ThermoFisher Scientific | D1306 |
| DMEM | Gibco - Life technologies | 11995-065 |
| Fetal Bovine Serum | Gibco - Life technologies | 12483-020 |
| Pen Strep (Penicillin Streptomycin) | Gibco - Life technologies | 15140-122 |
| Plasmocin | InvivoGen | ant-mpt |
| Attachment Factor 1X | Gibco - Life technologies | S-006-100 |
| Lipofectamine 2000 | ThermoFisher Scientific | 11668019 |
| Magnetofectamine O2 | OZ Biosciences | MTX2-0750 |

| | | |
|---|---|---|
| Phusion High-Fidelity DNA Polymerase | ThermoFisher Scientific | F530L |
| Western Lightning Plus ECL | PerkinElmer | NEL10400EA |
| Halt™ Protease Inhibitor Cocktail | ThermoFisher Scientific | 78425 |
| NEBExpress® Ni Spin Columns | New England Biolabs | S1427S |
| Slide-A-Lyzer™ MINI Dialysis Devices | ThermoFisher Scientific | 69570 |
| Pierce™ BCA Protein Assay Kit | ThermoFisher Scientific | 23227 |
| Recombinant *E. coli* MBP His Protein | Novus Biologicals | NBP2-22654 |
| CytoTox-Glo™ Cytotoxicity Assay | Promega | G9291 |
| NanoBit Protein:Protein Interaction System | Promega | N2014 |
| Nano-Glo Live Cell Assay System | Promega | N2012 |
| Recombinant DNA | | |
| Plasmid: SmBiT-PRKACA Control Vector | Promega | N2014 |
| Plasmid: LgBiT-PRKAR2A Control Vector | Promega | N2014 |
| Plasmid: NanoBiT Negative Control Vector | Promega | N2014 |
| Plasmid: pBiT-N-Lg-TDP-43 | This paper | N/A |
| Plasmid: pBiT-C-Lg-TDP-43 | This paper | N/A |
| Plasmid: pBiT-N-Lg-TDP-43$^{\Delta NLS}$ | This paper | N/A |
| Plasmid: pBiT-C-Lg-TDP-43$^{\Delta NLS}$ | This paper | N/A |
| Plasmid: pBiT-C-Lg-TDP-43$^{(1-366)}$ | This paper | N/A |
| Plasmid: pBiT-C-Lg-TDP-43$^{(1-274)}$ | This paper | N/A |
| Plasmid: pBiT-C-Lg-TDP-43$^{\Delta RRM-1-2}$ | This paper | N/A |
| Plasmid: pBiT-C-Lg-TDP-43$^{\Delta RRM-1}$ | This paper | N/A |
| Plasmid: pBiT-C-Lg-TDP-43$^{\Delta RRM-2}$ | This paper | N/A |
| Plasmid: pBiT-C-Lg-TDP-43$^{(1-192)}$ | This paper | N/A |
| Plasmid: pBiT-N-Sm-RGNEF | This paper | N/A |

| | | |
|---|---|---|
| Plasmid: pBiT-C-Sm-RGNEF | This paper | N/A |
| Plasmid: pBiT- N-Sm-NF242 | This paper | N/A |
| Plasmid: pBiT-C-Sm-NF242 | This paper | N/A |
| Plasmid: pQE30-TDP-43$^{1\text{-}261}$ | Emanuele Buratti (Italy) | N/A |
| Plasmid : pBAD-HisA-GST-TDP-43Cri | Emanuele Buratti (Italy) | N/A |
| Plasmid: pQE30-TDP-43-RRM1 | This paper | N/A |
| Plasmid: pQE30-TDP-43-RRM2 | This paper | N/A |
| Plasmid: pDEST566-RGNEF-275 | Murray Junop (Canada) | N/A |

#### Supplementary Table 2

| Insert | Backbone | Vector name | Protein product |
|---|---|---|---|
| TDP-43 | pBiT1.1-N [TK LgBiT] | pBiT-N-Lg-TDP-43 | Lg-TDP-43 |
| | pBiT1.1-C [TK LgBiT] | pBiT-C-Lg-TDP-43 | TDP-43-Lg |
| TDP-43-ΔNLS | pBiT1.1-N [TK LgBiT] | pBiT-N-Lg-TDP-43-ΔNLS | Lg-TDP-43-ΔNLS |
| | pBiT1.1-C [TK LgBiT] | pBiT-C-Lg-TDP-43-ΔNLS | TDP-43-ΔNLS-Lg |
| RGNEF | pBiT2.1-N [TK SmBiT] | pBiT-N-Sm-RGNEF | Sm-RGNEF |
| | pBiT2.1-C [TK SmBiT] | pBiT-C-Sm-RGNEF | RGNEF-Sm |
| NF242 | pBiT2.1-N [TK SmBiT] | pBiT- N-Sm-NF242 | Sm-NF242 |
| | pBiT2.1-C [TK SmBiT] | pBiT-C-Sm-NF242 | NF242-Sm |

#### Supplementary Table 3

(Fly lines from stock centers)

| Name | Stock# | Company | Chr. | Expression |
|---|---|---|---|---|
| GMR | 1104 | Bloomington | 2 | Eye* |
| D42 | 8816 | Bloomington | 3 | Motor Neuron |
| elav | 458 | Bloomington | 1 | Pan-Neuronal† |
| w- | 60000 | Vienna Drosophila Resource Center | 1 | --- |
| GFP; Dr/Sb | 60292 | Bloomington | 2 | --- |
| C9-36R | 58688 | Bloomington | 2 | --- |

* = Provides strong expression in all cells behind the morphogenetic furrow.

† = Begins expression at stage 12 of embryonic development.

#### Supplementary Table 4

| Gene expressed | Driver | Genotype | Name in this manuscript |
|---|---|---|---|
| RGNEF | elav | $\frac{w[1118]}{elav-GAL4};\frac{UAS-RGNEF.myc,mw+}{+};\frac{+}{+}$ | *elav>RGNEF* |
| NF242 | GMR | $\frac{w[1118]}{+};\frac{UAS-flag.NF242,mw+}{GMR-GAL4};\frac{+}{+}$ | *GMR>NF242* |
| GFP + TDP-43 | GMR | $\frac{w[1118]}{+};\frac{UAS-2xEGFP}{GMR-GAL4};\frac{UAS-hTDP43,mw+}{+}$ | *GMR>GFP;TDP-43$^{wt}$* |
| | elav | $\frac{w[1118]}{elav-GAL4};\frac{UAS-2xEGFP}{+};\frac{UAS-hTDP43,mw+}{+}$ | *elav>GFP;TDP-43$^{wt}$* |
| | D42 | $\frac{w[1118]}{+};\frac{UAS-2xEGFP}{+};\frac{UAS-hTDP43,mw+}{D42-GAL4}$ | *D42>GFP;TDP-43$^{wt}$* |
| RGNEF + TDP-43 | GMR | $\frac{w[1118]}{+};\frac{UAS-RGNEF.myc,mw+}{GMR-GAL4};\frac{UAS-hTDP43,mw+}{+}$ | *GMR>RGNEF;TDP-43$^{wt}$* |
| | elav | $\frac{w[1118]}{elav-GAL4};\frac{UAS-RGNEF.myc,mw+}{+};\frac{UAS-hTDP43,mw+}{+}$ | *elav>RGNEF;TDP-43$^{wt}$* |
| | D42 | $\frac{w[1118]}{+};\frac{UAS-RGNEF.myc,mw+}{+};\frac{UAS-hTDP43,mw+}{D42-GAL4}$ | *D42>RGNEF;TDP-43$^{wt}$* |
| NF242 + TDP-43 | GMR | $\frac{w[1118]}{+};\frac{UAS-flag.NF242,mw+}{GMR-GAL4};\frac{UAS-hTDP43,mw+}{+}$ | *GMR>NF242;TDP-43$^{wt}$* |
| | elav | $\frac{w[1118]}{elav-GAL4};\frac{UAS-flag.NF242,mw+}{+};\frac{UAS-hTDP43,mw+}{+}$ | *elav>NF242;TDP-43$^{wt}$* |
| | D42 | $\frac{w[1118]}{+};\frac{UAS-flag.NF242,mw+}{+};\frac{UAS-hTDP43,mw+}{D42-GAL4}$ | *D42>NF242;TDP-43$^{wt}$* |

#### Supplementary Table 5

| Primer name | Sequence | Amplicon size |
|---|---|---|
| TDP-43-F | 5’ GGACTTGATCATTAAAGGAATCAGCGTTC 3’ | 447 bp |
| TDP-43-R | 5’ CTGCCCGACCCTGCATTGGATG 3’ | |
| RGNEF-F | 5’ GCCCCGAGGTAATGGAACTTAATCG 3’ | 550 bp |
| RGNEF-R | 5’ TAAACAATATTTTCTTTGGCTCCATCTCCAGT 3’ | |
| NF242-F | 5’ ATGACAAGATGGAGTTGAGCTGCAGCGAAG 3’ | 734 bp |
| NF42-R | 5’ GTAATGCAAGGAGGCTTCTTCACTG 3’ | |
| GFP-F | 5’ CCACCCTCGTGACCACCCTGA 3’ | 474 bp |
| GFP-R | 5’ CGCGCTTCTCGTTGGGGTCTT 3’ | |

### Supplementary Table 6

(Antibody dilutions)

| Antibody | Dilution |
| --- | --- |
| Anti-flag (goat) | 1/200 |
| Anti-GFP (rabbit) | 1/400 |
| Anti-TDP-43 (rabbit) | 1/250 (IF) 1/4000 (WB) |
| Anti-TDP-43 (mouse) | 1/250 |
| Anti-Iba1 (rabbit) | 1/100 |
| Anti-GFAP (mouse) | 1/100 |
| Anti-goat Alexa 488 (donkey) | 1/1,000 |
| Anti-rabbit Alexa 555 (donkey) | 1/1,000 |
| Anti-mouse Alexa 555 (donkey) | 1/1,000 |
| Anti-rabbit Alexa 594 (donkey) | 1/1,000 |
| Anti-rabbit Alexa 680 (donkey) | 1/1,000 |

### Supplementary Methods

#### Protein Purification

For purification of His-GST-TDP-43$^{wt}$, transformed *E. coli* BL21 (DE3) cells were grown in LB medium. The expression was induced by adding 0.2% L-Arabinose and the culture was grown for an additional 4 hours at 37 ºC. The cells were harvested by centrifugation and resuspended in lysis buffer (20 mM Tris-HCl, pH 7.5, 500 mM NaCl, 20 mM Imidazole, 10% Glycerol, 1 mM PMSF, and 1x Protease Inhibitor Cocktail). The cells were lysed using EmulsiFlex-C3 (Avestin) and Bioruptor UCD-200 (Diagenode). The lysate was clarified by centrifugation and the supernatant was loaded onto a column packed with Ni-IDA resin (GE Life Sciences) pre-equilibrated with wash buffer (20 mM Tris-HCl, pH 7.5, 500 mM NaCl, 20 mM Imidazole, 10% Glycerol). The column was washed with wash buffer, and the bound protein was eluted with elution buffer (20 mM Tris-HCl, pH 7.5, 500 mM NaCl, 250 mM Imidazole, 10% Glycerol). The eluted protein was passed through a GPC column (Superdex 75 Increase 10/300 GL, GE Life Sciences) equilibrated with buffer (20 mM Tris-HCl, pH 7.5, 150 mM NaCl, 10% Glycerol). The major peak obtained in GPC was pooled and further concentrated using Amicon Ultra-15 centrifugal filters (Millipore) with a molecular weight cutoff of 10 kDa. The purified protein was dialyzed to 0.2 µm-filtered SPR running buffer (20 mM Hepes-7.4, 50 mM NaCl, 50 mM KCl, 0.5 mM MgCl2, 0.05% Tween-20, pH 7.4) using 10 kDa molecular weight cut off Slide-A-Lyzer™ MINI Dialysis Devices following manufacturer instructions immediately prior to SPR analysis.

For the purification of His-TDP-43$^{1\text{-}269}$, His-RRM-1, and His-RRM-2, M15 bacteria were grown in liquid LB containing ampicillin (Amp) and kanamycin (Kan) (ampicillin resistance encoded on the plasmid and kanamycin resistance encoded by the M15 strain of *E. coli*) at 37 °C overnight. For His-MBP-RGNEF$^{1\text{-}275}$ bacteria were grown in LB containing only Amp (BL21DE3T1R *E. coli*). The expression was induced with 1 mM isopropyl β-d-1-thiogalactopyranoside (IPTG) at 18°C for 16 hours. Induced cultures were collected, centrifuged, and then resuspended in cold lysis buffer containing 1x PBS, 1x Halt™ Protease Inhibitor Cocktail and 1 mg/ml lysozyme for TDP-43 proteins and 800 mM NaCl, 20 mM Tris-HCl pH 8.0, and 10% v/v glycerol for His-MBP-RGNEF$^{1\text{-}275}$. Pellets were incubated on ice for 30 minutes and lysed by sonication or French Press. Lysates containing His-TDP-43$^{1\text{-}269}$, His-RRM-1, and His-RRM-2 were cleared by centrifugation and applied to NEBExpress® Ni Spin

Columns following manufacturer directions. For His-MBP-RGNEF$^{1-275}$ Ni-IMAC was used on the ÄKTA Start Protein Purification System with a 5 mL HiTrap HP column (Cytiva Life Sciences, United States). Used buffers were: lysis buffer as described above, wash buffer (1x PBS, 300 mM NaCl, 5 mM imidazole for His-TDP-43$^{1-269}$, His-RRM-1, and His-RRM-2; and an imidazole gradient up to 800 mM NaCl, 300 mM imidazole, 20 mM Tris-HCl pH 8.0, and 10% v/v glycerol for His-MBP-RGNEF$^{1-275}$), and elution buffer (1x PBS, 1 mM DTT, 0.1% v/v Tween-20, 500 mM imidazole for His-TDP-43$^{1-269}$, His-RRM-1, and His-RRM-2; and 800 mM NaCl, 300 mM imidazole, 20 mM Tris-HCl pH 8.0, and 10% v/v glycerol for His-MBP-RGNEF$^{1-275}$). Purified proteins were visualized by SDS-PAGE. Eluted samples used for SPR analysis were incubated on ice and dialyzed to 0.2 μm-filtered SPR running buffer using 10 kDa molecular weight cut-off Slide-A-Lyzer™ MINI Dialysis Devices according to the manufacturer instructions immediately prior to SPR analysis.

Protein concentrations were determined using the BCA microplate assay and BSA standards according to the manufacturer directions.

#### Supplementary videos

**Supplementary video 1.** Video showing the phenotype of a rNLS8 mouse injected with AAV9/GFP after 3 weeks without Dox.

**Supplementary video 2.** Video showing the phenotype of a rNLS8 mouse injected with AAV9/GFP after 5 weeks without Dox.

**Supplementary video 3.** Video showing the phenotype of a rNLS8 mouse injected with AAV9/NF242 after 3 weeks without Dox.

**Supplementary video 4.** Video showing the phenotype of a rNLS8 mouse injected with AAV9/NF242 after 5 weeks without Dox.
